# Ultra-low photodamage three-photon microscopy assisted by neural network for monitoring regenerative myogenesis

**DOI:** 10.1101/2024.08.11.607469

**Authors:** Yifei Li, Keying Li, Mubin He, Chenlin Liang, Xin Xie, Jun Qian

## Abstract

Three-photon microscopy (3PM) enables high-resolution three-dimensional (3D) imaging in deeply situated and highly scattering biological specimens, facilitating precise characterization of biological morphology and cellular-level physiology *in vivo*. However, the use of fluorescent probes with relatively low three-photon absorption cross-sections necessitates high-peak-power lasers for excitation, which poses inherent risks of light-induced damage. Additionally, the low repetition frequency of these lasers prolongs scanning time per pixel, hampering imaging speed and exacerbating the potential for photodamage. Such limitations hinder the application of 3PM in studying vulnerable tissues, including muscle regeneration. To address this critical issue, we developed the Multi-Scale Attention Denoising Network (MSAD-Net), a precise and versatile denoising network suitable for diverse structures and varying noise levels. Our network enables the use of lower excitation power (1/4-1/2 of the common power) and shorter scanning time (1/6-1/4 of the common time) in 3PM while preserving image quality and tissue integrity. It achieves an impressive structural similarity index (SSIM) of up to 0.9932 and an incredibly fast inference time of just 80 milliseconds per frame which ensured both high fidelity and practicality for downstream applications. By utilizing MSAD-Net-assisted imaging, we comprehensively characterize the biological morphology and functionality of muscle regeneration processes through deep *in vivo* five-channel imaging under extremely low excitation power and short scanning time, while maintaining a high signal-to-background ratio (SBR) and excellent axial spatial resolution. Furthermore, we conducted high axial-resolution dynamic imaging of vascular microcirculation, macrophages, and ghost fibers. Our findings provide a deeper understanding of the mechanisms underlying muscle regeneration at the cellular and tissue levels.

## INTRODUCTION

Skeletal muscle possesses a remarkable ability to regenerate after damage from exercise or muscle disease, such as Duchenne Muscular Dystrophy (*1, 2*). Muscle regeneration is a highly orchestrated process that is primarily mediated by a population of resident muscle stem cells (MuSCs, also called satellite cells) (*3, 4*). Upon injury, activated satellite cells proliferate, differentiate, and fuse together to repair damaged myofibers (*2, 5, 6*). Therefore, precise control over satellite cells behavior is essential for proper regeneration and muscle homeostasis. Recent discoveries show that fate choices of satellite cell in regenerative myogenesis are dictated by a dynamically interactive microenvironment, which allows satellite cells to communicate with the numerous distinct cell types in their niche (*7, 8*). However, identifying and intervening the complex cell interactions has been hampered by an inability to accurately 3D visualize cell events and dynamics that occur within the regeneration.

In recent years, three-photon microscopy (3PM) has emerged as a powerful imaging tool, finding application in various tissues such as the mouse/monkey brain (*9–13*), adult zebrafish brain (*14*), mouse lymphoid tissues (*15*), mouse tumors (*16*) and mouse liver (*17–19*). Unlike the widely employed two-photon microscopy (2PM), 3PM utilizes longer-wavelength excitation laser beams, resulting in lower photon scattering and enhanced penetration capabilities. Additionally, it exhibits superior nonlinear optical confinement in highly scattering tissues, facilitating precise visualization of biological morphology and cellular-level physiology *in vivo*, particularly in densely labeled biological samples (*15, 19, 20*). 3PM presents a promising tool for investigating muscle tissue which contains dense fibers and abundant blood vessels. However, the use of fluorescent probes with relatively low three-photon absorption cross-sections necessitates high-peak-power lasers for excitation in 3PM. This heightened power requirement raises concerns regarding potential light-induced damage, such as photothermal damage and ionization disruption (*21–24*). Furthermore, the low repetition frequency associated with high-peak-power lasers in 3PM translates to prolonged scanning times during imaging. This reduced observation speed not only hampers imaging efficiency but also exacerbates the risk of photodamage, particularly in studying vulnerable biological samples, including regenerative myogenesis.

To mitigate the photodamage caused by 3PM, several optimization methods have been developed (*25*). Yildirim et al. achieved non-invasive measurements of the visual response area of the cerebral cortex neurons in awake mice by developing a pre-chirp system to compensate for pulse broadening in the microscope and designing a scan and tube lens integrated with the objective lens to reduce the aberration in the microscope (*26*). Li et al. employed an adaptive excitation light source and pre-scan in 3PM to select regions of interest (ROI), thus effectively minimized photodamage by reducing excitation outside the ROI (*27*). However, these optimizations require specialized complex optical devices, limiting accessibility for many biomedical researchers.

Recently, deep learning method has shown promise in improving imaging speed, signal-to-background ratio (SBR) and spatial resolution of various imaging techniques, including confocal microscopy, 2PM and super-resolution microscopy (*28–33*). For example, Dai’s group proposed several self-supervised-learning-based denoising frameworks, including DeepCAD (*29*) and DeepSeMi (*31*). They were capable of spatiotemporal enhancement of calcium imaging data in 2PM and recording organelle interactions in four colors at high frame rates across tens of thousands in super-resolution microscopy, respectively. In addition, Lu et al. also developed a supervised deep-denoising method to circumvent tradeoffs between the quality of the extracted calcium traces, imaging speed and laser power, for recovering complex neurite structures of *C. elegans* in confocal microscopy (*28*). To avoid the photodamage in 3PM, the implementation of short scanning time and low excitation power would result in a dramatic reduction of SBR (*26*). Deep learning techniques should be useful in denoising of 3PM images, which enables the use of lower laser power and shorter scanning time without compromising the SBR of the captured images.

Here, we developed the Multi-Scale Attention Denoising Network (MSAD-Net), a highly precise and versatile network suitable for small datasets. Our network enables the use of lower excitation power (1/4-1/2 of the usual power) and shorter scanning time (1/6-1/4 of the usual time) in 3PM while preserving image quality and the integrity of biological tissue. It achieves an impressive structural similarity index (SSIM) of up to 0.9932 when compared with three-photon fluorescence images captured under high-peak-power excitation and prolonged scanning periods. This ensures the fidelity and precision of the denoised images, effectively preventing the generation of spurious structures and preserving the integrity of the original biological features. Furthermore, our network demonstrates real-time (80 milliseconds per frame) denoising capability for diverse structures and varying levels of noise, ensuring its robustness and practicality in various applications. Using the high-fidelity MSAD-Net-assisted imaging, we generated a spatialtemporal map of MuSCs, macrophages, muscular blood vessels, and myofibers, highlighting the role of macrophages and blood vessels in guiding MuSC-mediated muscle regeneration. We then conducted high axial-resolution dynamic imaging of vascular microcirculation to elucidate the regulatory mechanisms of blood vessels on MuSCs. Furthermore, we performed multiple 3D imaging of macrophages under low excitation power and short scanning time and tracked the motion trajectory of macrophages. These studies significantly enhanced our understanding of the cell-cell interactions underlying muscle regeneration.

## RESULTS

### Three-photon fluorescence imaging of mouse muscle tissue

To enable deep *in vivo* observation of the tibialis anterior (TA) muscle, we first captured images of vascular endothelial cells labeled with GFP and MuSCs labeled with tdTomato using three-photon excitation (3PE) and two-photon excitation (2PE) microscopy (fig. S1). Three-photon fluorescence imaging revealed superior depth penetration (350 μm) and higher SBR compared to two-photon fluorescence imaging (Fig. 1A-B, fig. S2A). Y-Z plane images further demonstrated that the axial resolution of MuSCs under 3PE exceeded that of 2PE, particularly at greater depths. Even at shallower depths, adjacent MuSCs were more clearly resolved under 3PE than under 2PE (Fig. 1B, fig. S3A-F). SBRs of images under 3PE maintained high throughout 0-350 μm depth while SBR of images under 2PE were much lower, extremely at depths over 200 μm (Fig. 1C). Subsequently, we analyzed the cell morphology of MuSCs from both types of microscopic images (fig. S3G-H). Specifically, while the average bounding box dimensions in the x and y directions were similar (approximately 20-30 μm) for both imaging methods, the bounding box length in the z direction was notably larger in the two-photon fluorescence images (∼40 μm) than in the three-photon images (around 27 μm, Fig. 1D). This disparity indicated z-axis stretching in the MuSC images captured with 2PE, leading to inaccuracies in morphological extraction.

**Fig. 1.**
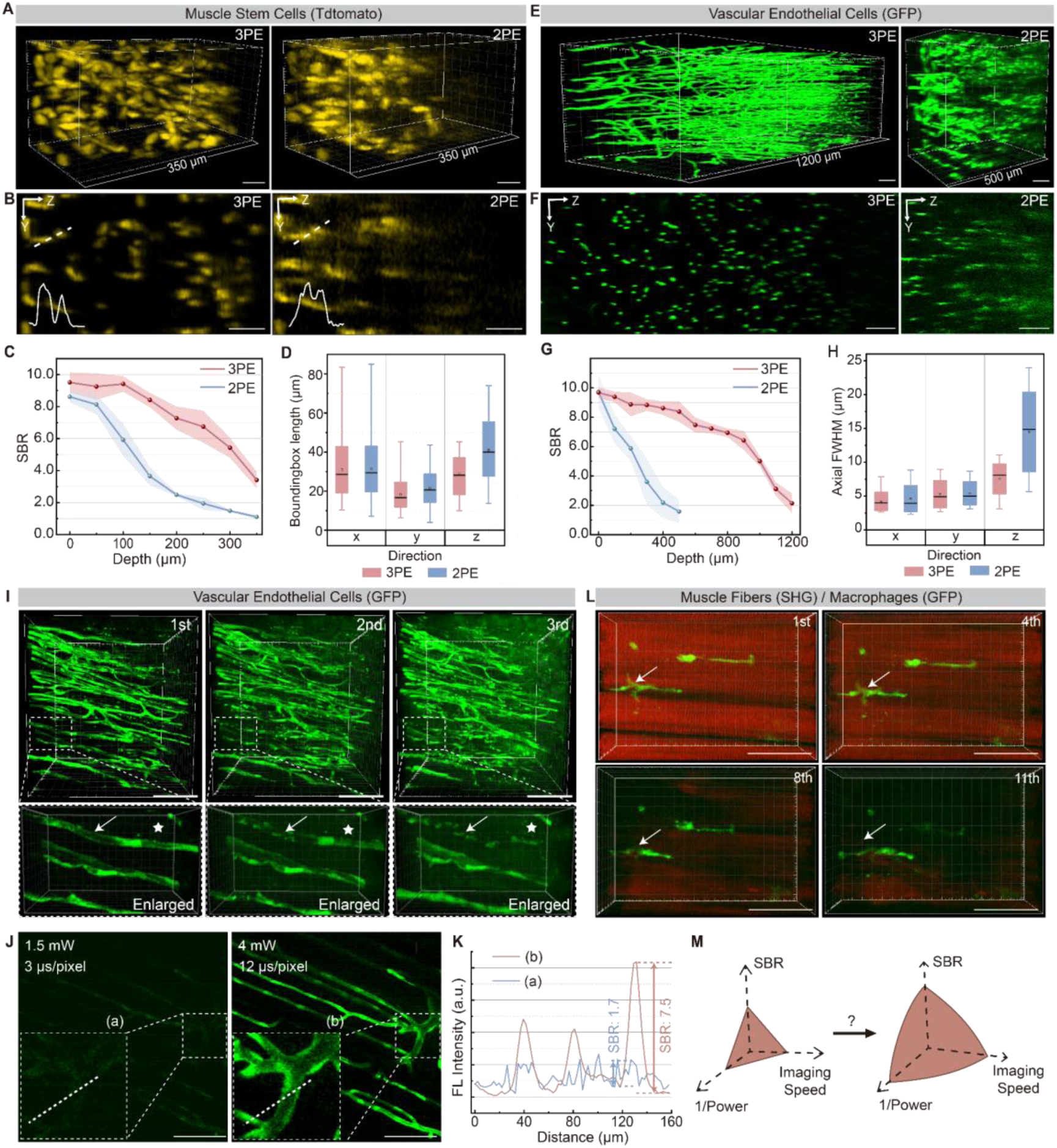
Three-photon microscopy enables deep imaging of mouse muscle tissue but causes tissue damage. (A) 3D reconstruction of three-photon and two-photon fluorescence microscopic images of MuSCs, which were labeled with tdTomato. Volume size: 200 μm ×200 μm ×350 μm. Scale bars: 50 μm. (B) Y-Z plane three-photon and two-photon fluorescence microscopic images of MuSCs. Bottom-left curves showed intensity along dotted lines in Y-Z plane images. Scale bars: 50 μm. (C) SBR of MuSCs fluorescence images as a function of imaging depth, derived from (a). (D) Bounding box lengths in X-Y-Z directions calculated from two-photon and three-photon fluorescence microscopic images of MuSCs shown in (A). (E) 3D reconstruction of three-photon (volume size: 400 μm ×400 μm ×1200 μm) and two-photon (volume size: 400 μm ×400 μm ×500 μm) fluorescence microscopic images of vascular endothelial cells, which were labeled with GFP. Scale bars: 100 μm. (F) Y-Z plane three-photon and two-photon fluorescence microscopic images of vascular endothelial cells. Scale bars: 100 μm. (G) SBR of vascular endothelial cells fluorescence images as a function of imaging depth, derived from (E). (H) Axial Full Width at Half Maximum (FWHM) in X-Y-Z directions calculated from two-photon and three-photon fluorescence microscopic images of vascular endothelial cells shown in (E). (I) Three-photon fluorescence microscopic images of vascular endothelial cells taken three times continuously (volume size: 400 μm ×400 μm ×350 μm). Below panels showed the enlarged structure of vascular endothelial cells (volume size: 120 μm ×72 μm ×50 μm). White arrows indicated the damaged vascular endothelial cells during continuous imaging. White pentagram stars indicated the stray spontaneous signal points during continuous imaging Scale bars: 100 μm. (J) Three-photon fluorescence microscopic images of vascular endothelial cells under two different imaging conditions (low excitation power and short scanning time vs normal excitation power and prolonged scanning time). Bottom-left panels showed the enlarged structure of the area within the dashed boxes. Scale bars: 100 μm. (K) Intensity profiles along dotted lines across vascular endothelial cells images in (J). The values of SBR indicated in the figures were calculated as maximum signal intensity divided by the background intensity according to the intensity profile. (L) Three-photon fluorescence microscopic images of macrophages and SHG microscopic images of fibers taken 11 times continuously (volume size: 170 μm ×120 μm ×64 μm). White arrows indicated the damaged macrophages during continuous imaging. Scale bars: 50 μm. (M) How to overcome the tradeoff between excitation power/imaging speed and SBR during three-photon fluorescence microscopic imaging.

Regarding vascular endothelial cells, 3D images captured using 3PE consistently exhibited higher SBR across the entire depth range (Fig. 1E-F and fig. S2B). In contrast, images obtained with 2PE showed a notable decrease in SBR with increasing depth, as depicted in Fig. 1G. Furthermore, axial vessel diameters observed in 2PE images appeared larger than those in 3PE images, suggesting inferior axial confinement with 2PE (Fig. 1H). Three-photon fluorescence imaging of vascular endothelial cells revealed well-organized blood vessels with consistently high SBR at various depths (fig. S4, movie. S1). Collectively, these findings underscore the advantages of 3PM over 2PM in terms of imaging depth, SBR, and axial resolution.

To explore muscle injury-induced regeneration through cellular-level imaging, avoiding additional tissue damage is crucial. We began by analyzing SBRs of three-photon fluorescence microscopic images under varied excitation powers and scanning durations (fig. S5-6). Imaging conditions, with laser power exceeding 8 mW and scanning times over 8 μs/pixel, facilitated clear visualization of macrophages (SBR > 4). Conversely, conditions with laser power below 4 mW and scanning times below 6 μs/pixel posed challenges in discerning macrophage features (SBR < 2). Figure 1i illustrates three-photon fluorescence microscopic images of vascular endothelial cells with sufficient SBR at depths ranging from 0 to 350 μm, at 1 day post injury (1 dpi). Reduced imaging depth was observed due to increased photon scattering in injured muscle tissue (fig. S7). Subsequent microscopic imaging on the same day led to noticeable light-induced damage: endothelial cells appeared deformed and irregular, as depicted by white arrows in Fig. 1I. Additionally, stray spontaneous signals emerged outside cells, denoted by white pentagram stars in Fig. 1I. Our findings indicated that 3PM induced photodamage to muscle tissues when ensuring adequate image SBR. Decreasing excitation power and scanning time to prevent photodamage resulted in less distinct endothelial cell structures (Fig. 1J), with SBR declining from 7.5 to 1.7 (Fig. 1K). To assess the broader impact of photodamage from 3PM on muscle tissue structures, we conducted continuous three-photon fluorescence imaging of macrophages and second harmonic generation (SHG) imaging of fibers. Results revealed weakening SHG signals with prolonged imaging (Fig. 1l, Supplementary Video 2). After 8-11 repeated imaging sessions, fibers were fragmented, accompanied with notable changes in macrophage morphology (white arrows). Thus, balancing photodamage with imaging quality is critical when conducting three-photon fluorescence imaging of muscle tissue. Efforts should focus on developing methodologies that achieve high SBR image with reduced excitation power and scanning time (Fig. 1M).

### A high-precision neural network for denoising images of 3PM

To address the tradeoff between photodamage and imaging quality in 3PM, we developed MSAD-Net, a highly precise and versatile network suitable for small datasets (fig. S8). To ensure the authenticity and generalizability of the output images, we propose an improved UNet model based on Transfomer and the Convolutional Block Attention Module (CBAM) attention mechanism. By introducing Transformer encoders and decoders, the model captures global contextual information and facilitates long-range information propagation, thereby enhancing the performance and generalization capabilities of the model. Additionally, we incorporated CBAM to replace the skip connection, enhancing the model’s receptive field and discriminative ability. Furthermore, we designed a hybrid loss function that combines mean square error (MSE) and multi-scale structural similarity index (MS-SSIM) to achieve higher accuracy (fig. S9). Moreover, we used two-dimensional (2D) images instead of 3D stacks to simplify the training steps and improve training speed, achieving an average inference time of 80 milliseconds per frame. Furthermore, our model has been rigorously tested and applied to different biological structures, showcasing its remarkable interpretability and robustness.

To evaluate the performance and robustness of our approach, we assessed the denoising ability of the network using signal characteristics from different structures of tibia anterior (TA) muscle tissue. The training process of the MSAD-Net is illustrated in Fig. 2A. Initially, we captured three-photon fluorescence images of TA muscle under normal excitation power and prolonged scanning time conditions to obtain high SBR. We then reduced the excitation power and scanning time to acquire images with low SBR corresponding to those with high SBR. Pairs of image data from different samples and structures were used as input to train the network. During the process, one-to-one corresponding training set images were captured on a fixed static sample to avoid the complex image pre-registration step across image pairs before training the network. The trained network was then applied to low SBR images (noisy images) to recover high SBR images. As shown in the upper-left panel of Fig. 2B-D, 3PM with low excitation power and short scanning time resulted in low SBR, making vascular endothelial cells, MuSCs, and macrophages indistinguishable from the background. Undistinguishable structures pose challenges for downstream analysis, such as morphology and dynamics tracking. By utilizing the MSAD-Net for denoising, we achieved significant restoration of various biological structures from the initially noisy images. The intensity analysis revealed that the image SBRs of vascular endothelial cells, MuSCs and macrophages increased from 1.7, 2.0, and 2.1 to 14.5, 27.7, and 30.2, respectively, after denoised by the MSAD-Net (Fig. 2B-D).

**Fig. 2.**
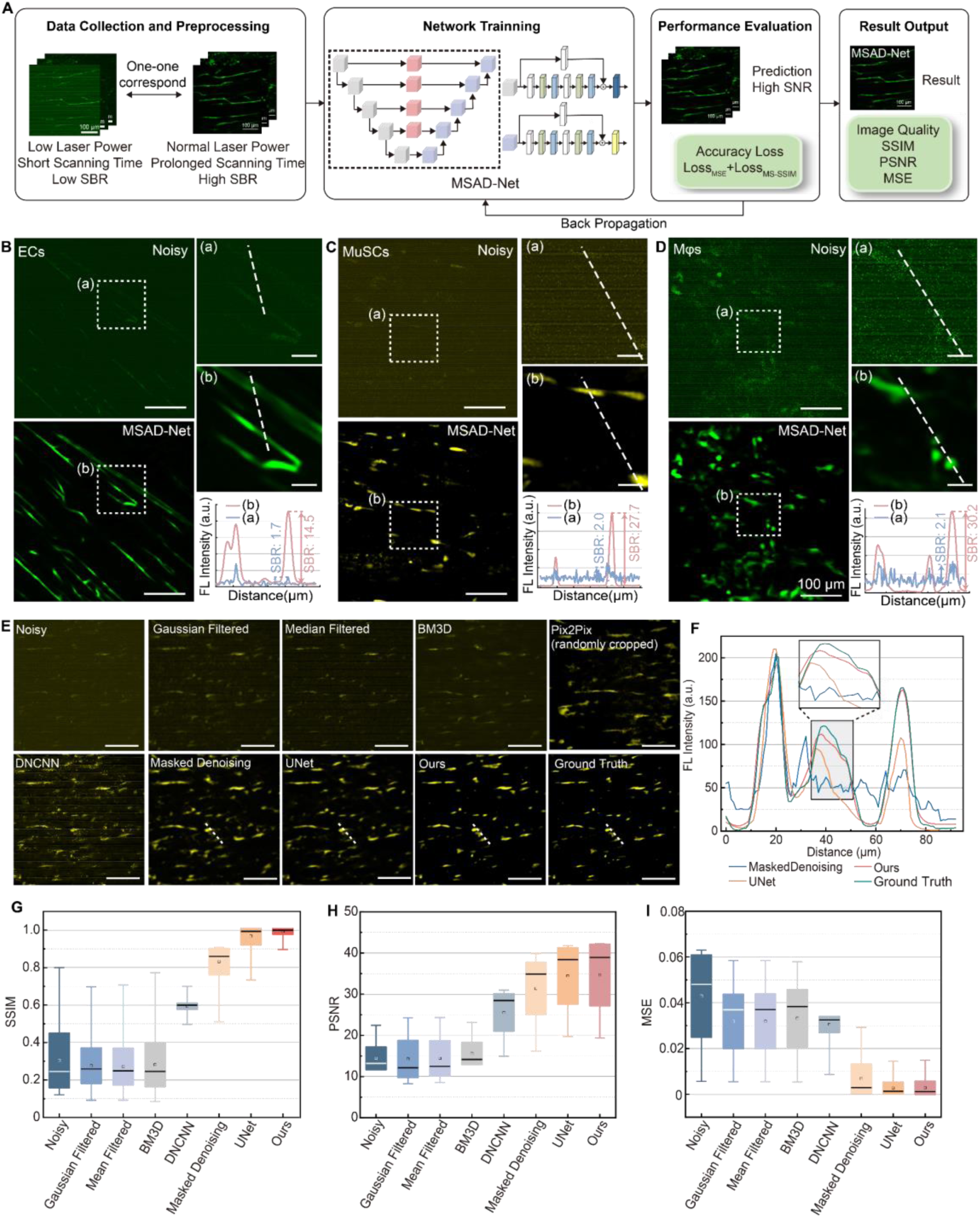
Three-photon fluorescence microscopic images denoised by the high-precision MSAD-Net. (A) Overview of the Multi-Scale Attention Denoising Network (MSAD-Net) to recover images of high SBR from those of low SBR. (B-D) Typical noisy images and corresponding denoised images of (B) vascular endothelial cells (ECs), (C) MuSCs, and (D) macrophages (Mφs) (scale bar: 100 μm). Top-left panels showed the noisy images. Bottom-left panels showed the images denoised by the MSAD-Net. Top-right panels showed insets ‘(a)’ and ‘(b)’ in noisy and denoised images (scale bar: 20 μm). Bottom-right panels showed intensity along dotted lines in noisy and denoised images in top-right panels. The values of SBR indicated in the figures were calculated as maximum signal intensity divided by the background intensity according to the intensity profile. (E) The original noisy image and corresponding denoised output images of MuSCs by various methods, including Gaussian filter, Median filter, BM3D, Pix2Pix, DNCNN, Masked Denoising, UNet and MSAD-Net. The image with high SBR was set as Ground Truth for comparison. Imaging depth: 150 μm. Imaging condition: 6 mW excitation power and 3 μs/pixel scanning time for noisy imaging while 15 mW excitation power and 12 μs/pixel scanning time for Ground Truth imaging. Scale bar: 100 μm. (F) Intensity profiles along the white dashed lines across MuSCs in (E). Insert was the enlarged intensity profiles in rectangular box. Three deep learning networks (Masked Denoising, UNet and MSAD-Net) with relatively good denoising effects were selected for comparison. (G-I) Comparison of (G) SSIM, (H) PSNR, and (I) MSE between ground-truth images with high SBR and noisy images, as well as denoised output images by various methods (including Gaussian filter, Median filter, BM3D, Pix2Pix, DNCNN, Masked Denoising, UNet and MSAD-Net, n = 340 images). Box center indicated median, box edges indicated standard deviation, and whiskers indicated 5th and 95th percentile.

To assess the superior ability of MSAD-Net to improve SBRs of images, we compared its performance with other approaches. Representative images demonstrate that simple denoising methods such as Gaussian and Median filtering, as well as advanced non-deep learning-based methods like BM3D, only partially reduced the background noise, resulting in persistent electrical noise and insufficient signal enhancement (Fig. 2E, fig. S10). We compared the image restoration capabilities of several deep learning networks, including Pix2pix, DNCNN, Masked Denoising, UNet, and our proposed network, using a dataset that included corresponding images with high and low SBRs under 3PE. Among these networks, the Pix2pix method randomly crops images and its denoising effect is generally limited, resulting in suboptimal restoration of cell morphology. DNCNN is trained by adding Gaussian noise to ground truth images and using both simulated noisy and high SBR images for training. However, this approach fails to accurately simulate the noise present in the imaging process. Although it can partially remove Gaussian noise, the overall performance remains unsatisfactory. On the other hand, the Masked Denoising network, UNet, and our proposed network exhibited relatively good denoising capabilities. Later, we compared the denoising effects of the three aforementioned deep learning methods. It is evident that our network produces restored images that are closer to the ground truth compared to the other networks based on the intensity profiles shown in Fig. 2F. Furthermore, we conducted additional performance comparisons among different methods based on SSIM, PSNR (peak signal-to-noise ratio), and MSE (mean squared error). Contrary to the other methods, our MSAD-Net exhibited exceptional denoising performance on low-noise images, specifically in preserving intricate details such as blood vessel and cell morphology. The PSNR value of our network reached approximately 34.7, indicating high-quality denoising. Additionally, the average SSIM value was notably high at 0.9932, as illustrated in Fig. 2G-H and table S1. Moreover, the MSE value was approximately 0.0028, further supporting the superior performance of our network (Fig. 2I and table S1). Overall, the denoised images produced by MSAD-Net were closer to the ground truth images and exhibited the strongest generalization ability.

### Spatial-temporal map of macrophages, MuSCs, muscular blood vessels, and myofibers during muscle regeneration

Next, we characterized the spatial-temporal properties of MuSCs and their microenvironment during regenerative myogenesis using the MSAD-Net. To visualize these components *in vivo*, we generated double-fluorescence labeled mice, where macrophages expressed GFP and MuSCs were marked with tdTomato (Fig. 3A). Additionally, we labeled the blood vessels via the injection of DCBT NPs, which have a large three-photon absorption cross-section, ensuring a great imaging depth(*12, 34*). Furthermore, SHG and third harmonic generation (THG) signals were produced from fibers and fiber membranes, respectively(*35, 36*). However, the fluorescence signals of GFP were significantly low under the femtosecond (fs) excitation wavelength of 1550 nm, while the fluorescence of tdTomato and SHG of fibers experienced crosstalk under 1300 nm excitation, making it impossible to image these structures using a single fs excitation wavelength. Hence, we performed four sets of deep dual-channel imaging of the muscle tissue using different fs excitation wavelengths, as explained in the methods, while employing low excitation power and short scanning time to minimize photodamage. Later, we merged the four sets of dual-channel images to obtain a five-channel muscle tissue imaging.

**Fig. 3.**
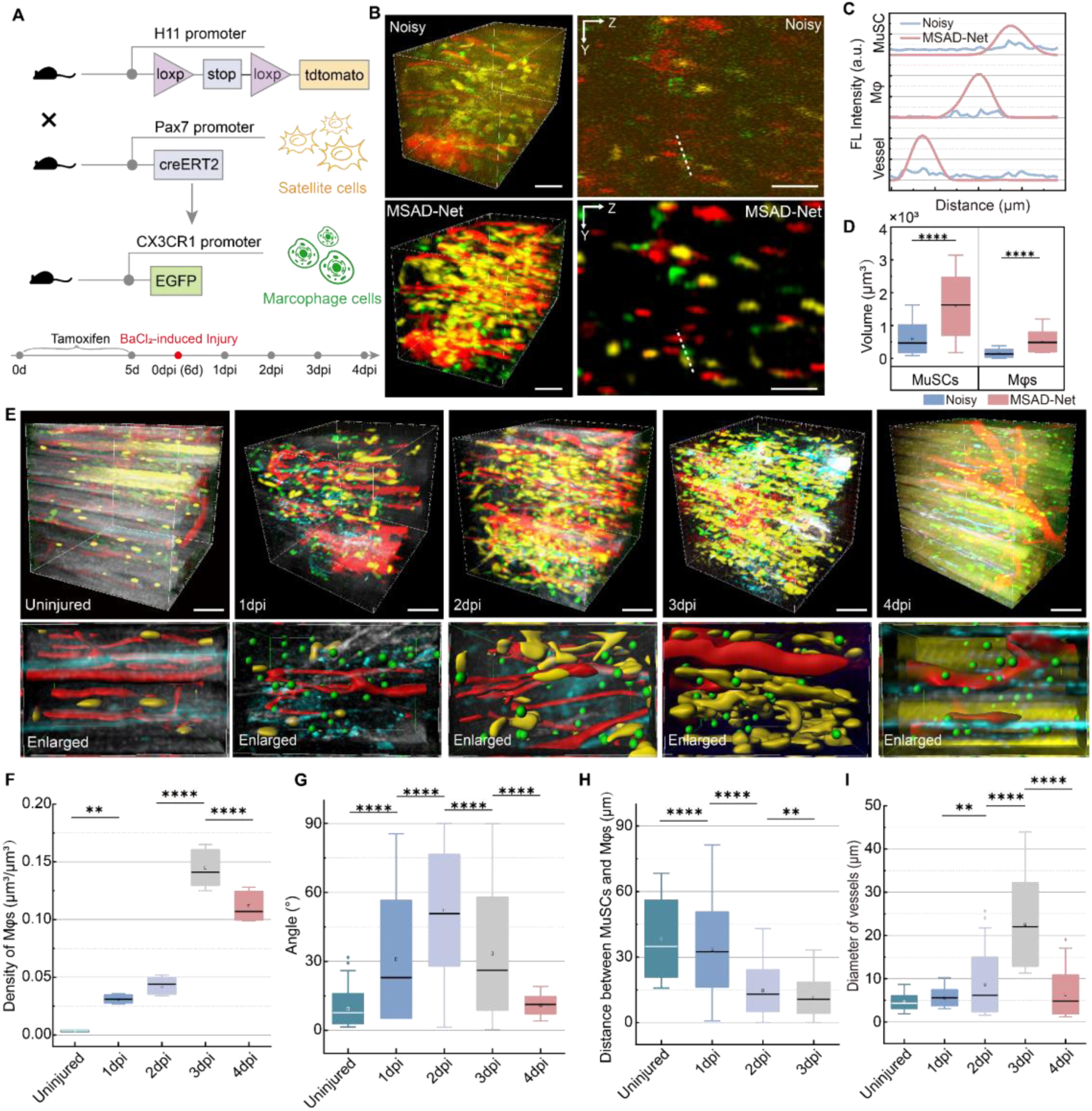
Five-channel microscopic imaging during muscle regeneration, including three-photon fluorescence imaging of macrophages, MuSCs and muscular blood vessels, SHG imaging of fibers, and THG imaging of fiber membranes, denoised by the MSAD-Net. (A) Schematic diagram illustrating the process of generating a targeting mouse hybrid model and the procedure of introducing mouse muscle injury model. H11-CAG-tdTomato mice were crossed with Pax7CreERT2/+ mice to generate mice where MuSCs were labeled with tdTomato. Subsequently these mice were crossed with CX3CR1-GFP mice to generate double-fluorescence labeled mice, where macrophages expressed GFP and MuSCs were labeled with tdTomato. Details of mouse muscle injury procedure were described in “Methods” part. (B) Three-channel three-photon fluorescence microscopic images of macrophages, MuSCs and muscle blood vessels before and after denoised by the MSAD-Net. The left panels displayed 3D structure images (volume size: 190 μm × 190 μm × 400 μm), while the right panels showed Y-Z plane images. Green represents macrophages, yellow represents MuSCs, red represents muscular blood vessels. Scale bar: 50 μm. (C) Intensity profiles along the dotted lines in the three-channel three-photon fluorescence microscopic images in (B). The identification of macrophages, MuSCs, and muscle blood vessels were significantly enhanced in the images denoised by the MSAD-Net. (D) Extracted volume of macrophages and MuSCs based on noisy images and denoised images (by the MSAD-Net). The box center indicates the median, the box edges represent the standard deviation, and the whiskers indicate the 5th and 95th percentiles. (E) Five-channel microscopic muscle images, including three-photon fluorescence microscopic images of macrophages, MuSCs and muscular blood vessels, SHG images of fibers, and THG images of fiber membranes, in uninjured mice and mice at 1-4 day(s) post-injury (1dpi-4dpi). Blue represents fiber membranes, green represents macrophages, yellow represents MuSCs, red represents muscular blood vessels, and gray represents fibers (details described in “Methods” part). The lower panels displayed enlarged five-channel images. MuSCs and muscular blood vessels were the morphological extraction results using “Surface” functionalities in the Imaris. Macrophages were the location results using “Spots” functionalities in the Imaris. Volume size: 340 μm × 340 μm × 400 μm. Scale bar: 100 μm. (F) Variation in the density of macrophages with injury time. (G) Variation in the angle between the long axis of the extracted MuSCs/regenerative muscle fibers and the direction of SHG signals with injury time. (H) Variation in the distance between MuSCs and macrophages with injury time. (I) Variation in the diameter of muscular blood vessels with injury time (calculated by DCBT NPs’ signals). Data in (F-I) were analyzed using one-way ANOVA with Tukey’s multiple comparison test. Error bars represent the standard deviation. *P < 0.05; **P < 0.01; ***P < 0.001; ****P < 0.0001.

The SBRs of three-photon fluorescence microscopic images of macrophages, MuSCs, and blood vessels were enhanced by the MSAD-Net. Fig. 3B illustrates a typical three-channel 3D image of muscle tissue after injury, acquired with low excitation power and short scanning time. Through the MSAD-Net, previously indistinguishable fluorescent structures in the input 3D image were restored. The denoised images exhibited an improved SBR and much cleaner structures compared to the original noisy images (Fig. 3B-C, fig. S11-14). Furthermore, we conducted morphology extraction using a watershed segmentation algorithm from a 3D input image. Comparing the segmentation results obtained from the noisy 3D images, the segmentation derived from the denoised 3D images identified an additional 57 MuSCs and 65 macrophages (fig. S15). Moreover, the volume of the cell morphology extracted using denoised 3D images was larger (Fig. 3D), enabling a more accurate obtain of cell signals.

The imaging results revealed that the uninjured TA muscle consisted of densely packed myofibers, as indicated by the SHG signals (Fig. 3E, uninjured). MuSCs were primarily located at the edges of the myofibers in a resting state, while macrophages were rarely detected at this stage. Following injury, the original myofibers gradually decomposed, accompanied by the disintegration of blood vessels, as well as the activation and proliferation of MuSCs and the recruitment of macrophages (Fig. 3E, 1-3dpi, Fig. 3F, Supplementary Fig. 16). The angle between the long axis of the extracted MuSCs and the direction of the SHG signal increased (Fig. 3G), suggesting that MuSCs proliferated in response to stimuli in a manner that deviated gradually from the direction of the myofibers. During this process, we observed an increasing proximity of macrophages to MuSCs (Fig. 3H) and an enlargement of blood vessel diameter (Fig. 3I). These features were more pronounced at 3 dpi when MuSCs started to fuse to form initial multinucleated myofibers. At 4 dpi, regenerative myofibers and blood vessels continued to grow accompanied with a decrease in macrophage density (Fig. 3F). However, SHG signals were almost undetectable at this time, consistent with the lack of collagen in early myotubes. In summary, we generated a spatialtemporal map of MuSCs, macrophages, blood vessels, and myotubes, highlighting the potential effects of macrophages and blood vessels in early regenerative myogenesis.

### Dynamic tracking of blood microcirculation during muscle regeneration

To explore the potential effects of blood vessels on MuSCs, we injected DCBT NPs into mice with endothelial cell-specific GFP reported to obtain a comprehensive dynamic map of microcirculation. Under 1300 nm fs excitation, dual-channel three-photon fluorescence microscopic imaging of endothelial cells (labeled with GFP) and blood microcirculation (stained with DCBT NPs) revealed the localization of endothelial cells at the edge of blood vessels, while the fluorescence signals of DCBT NPs were predominantly observed within the blood vessels (Fig. 4A-B). This distinction facilitated the differentiation of blood microcirculation from the vascular structure composed of endothelial cells. To minimize photodamage and restore the true signals of endothelial cells and blood vessels during continuous imaging, we employed low excitation power and short scanning time. The obtained images were then denoised using the MSAD-Net. As expected, the MSAD-Net-processed dual-channel images exhibited improved identification in defining the spatial positioning and structure of blood and vessel edges (Fig. 4C).

**Fig. 4.**
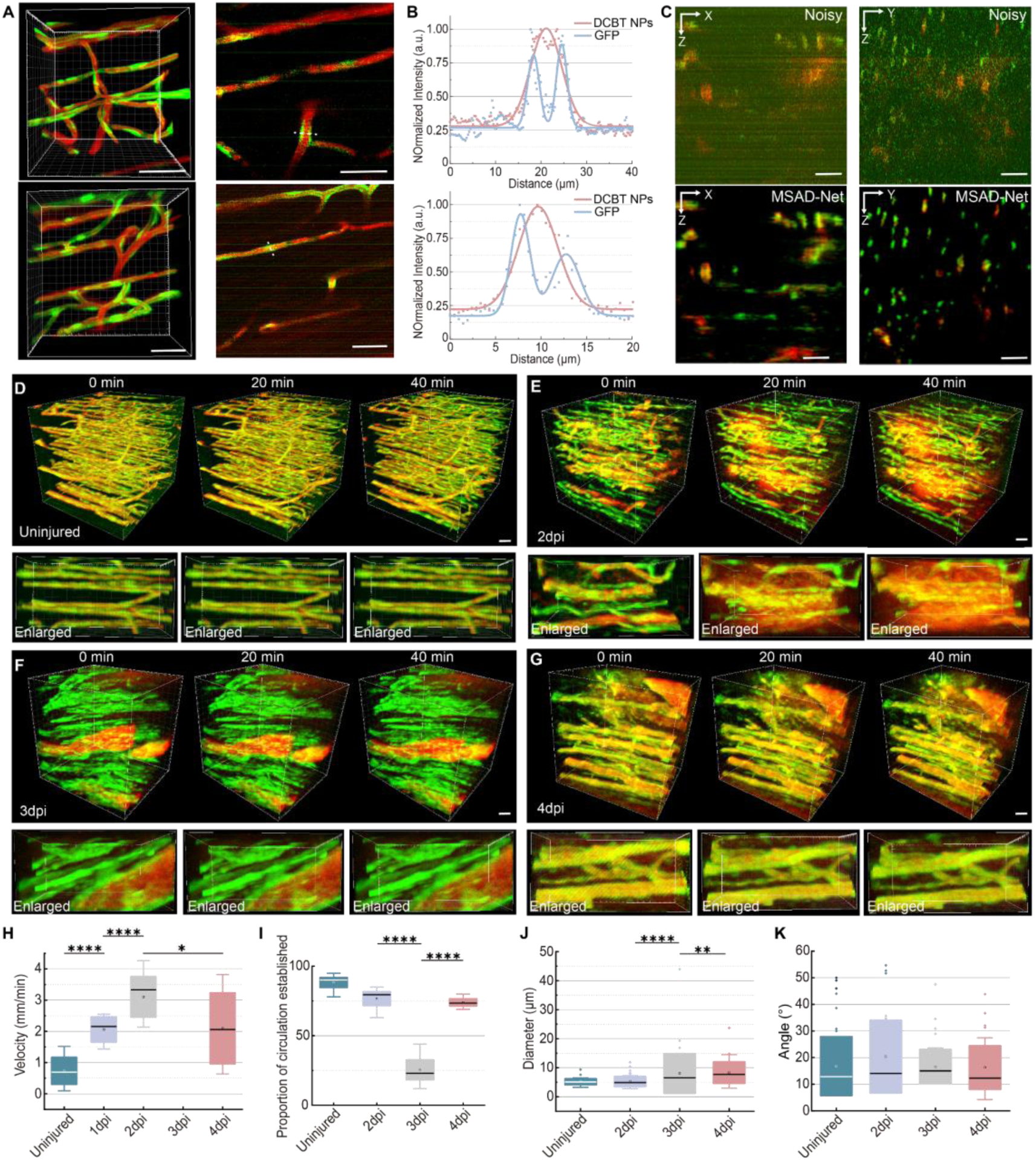
Dual-color three-photon fluorescence microscopic images of vascular endothelial cells and blood microcirculation during muscle regeneration, after denoised by the MSAD-Net. (A) Dual-channel three-photon fluorescence microscopic imaging of vascular endothelial cells (labeled by GFP) and blood microcirculation (labeled by DCBT NPs) under 4× zoom of a 25× objective view. The left panels showed two examples of 3D structure images, while the right panels displayed the corresponding X-Y plane images. Scale Bar: 10 μm. (B) Intensity profiles along the white dashed lines in the dual-channel images of panel (A). The signal of blood microcirculation was observed to be distributed in the middle of the endothelial cell signal. (C) Dual-channel X-Z and Y-Z three-photon fluorescence microscopic imaging of vascular endothelial cells and blood microcirculation, before and after denoised by the MSAD-Net. Scale bar: 50 μm. (D-G) Continuous dual-channel three-photon fluorescence microscopic imaging of vascular endothelial cells and blood microcirculation at 0, 2, 3 and 4 day(s) post-injury (Uninjured, 2dpi, 3dpi and 4dpi). The bottom panels showed locally enlarged 3D structure images. (H) Variation in the blood flow velocity of small blood vessels with a diameter less than 10 μm with injury time. (I) Variation in the proportion of circulation established with injury time. (J) Variation in the diameter of muscular blood vessels with injury time (calculated by vascular endothelial cells’ signals). (K) Variation in the angle between the direction of vascular endothelial cells and the original muscle fiber alignment with injury time. Data in (H-K) were analyzed using one-way ANOVA with Tukey’s multiple comparison test. Error bars represent the standard deviation. *P < 0.05; **P < 0.01; ***P < 0.001; ****P < 0.0001.

Under normal physiological condition, blood vessels maintained a relatively low blood flow velocity as shown by continuous three-photon fluorescence microscopic imaging (Fig. 4H). The unchanged DCBT NPs and GFP signals support the feasibility of our approach (Fig. 4D). We then monitored the dynamics of blood microcirculation during muscle repair (Fig. 4E-G). Interestingly, while the vasculature structure were disrupted upon damage, small blood vessels with a diameter less than 10 μm showed an increase in flow velocity (Fig. 4H, Supplementary Fig. 17). Additionally, a certain proportion of vessels did not exhibit DCBT NPs signals, indicating microcirculatory disturbances (Fig. 4I). Re-imaging the same locations after 20 minutes identified signs of DCBT NPs leakage, which was worsened after 40 minutes. At 3 dpi, only a small portion of vessels displayed DCBT NPs signals, implicating poor vascular connectivity at this stage (Fig. 4I). This hindered the establishment of circulation and resulted in blood leakage. Furthermore, an increase in the diameter of established vessels was observed, indicating a state of vessel growth (Fig. 3I, Fig. 4J). However, despite the increased vessel diameter, the blood flow velocity did not decrease but remained at a relatively high level, suggesting that the vessels may be experiencing higher pressure (Fig. 4H). At 4 dpi, the blood flow decreased, and DCBT NPs signals were detectable in most vessels, indicating a gradual restoration of microcirculation (Fig. 4G). At this stage, the formation of myofibers could be detected (Fig. 3E). Additionally, no significant alteration was observed in the orientation of endothelial cells during the injury process (Fig. 4K). Notably, previous studies proposed a “ghost fibers”, which guided the direction of newly formed myofibers during muscle repair (*36, 37*). Our results further suggested that endothelial cells also play a guiding role similar to “ghost fibers”.

In summary, through short-term continuous observations, we discovered that upon injury, vascular microcirculations were destroyed and permeability was increased, potentially leading to the release of inflammatory factors and the establishment of an inflammatory microenvironment.

### Dynamic 3D tracking map of macrophages during muscle regeneration

Next, we monitored macrophage movement during early regenerative myogenesis, aiming to create a dynamic migration map of these cells. To avoid photodamage caused by multiple-round 3D imaging, we employed low excitation power and short scanning time. We then applied the MSAD-Net to improve the SBR of images, and identified macrophage (Fig. 5A). As such, we successfully tracked the movement of macrophages under low excitation power and short scanning time, and delineated their motion trajectory (Fig. 5B).

**Fig. 5.**
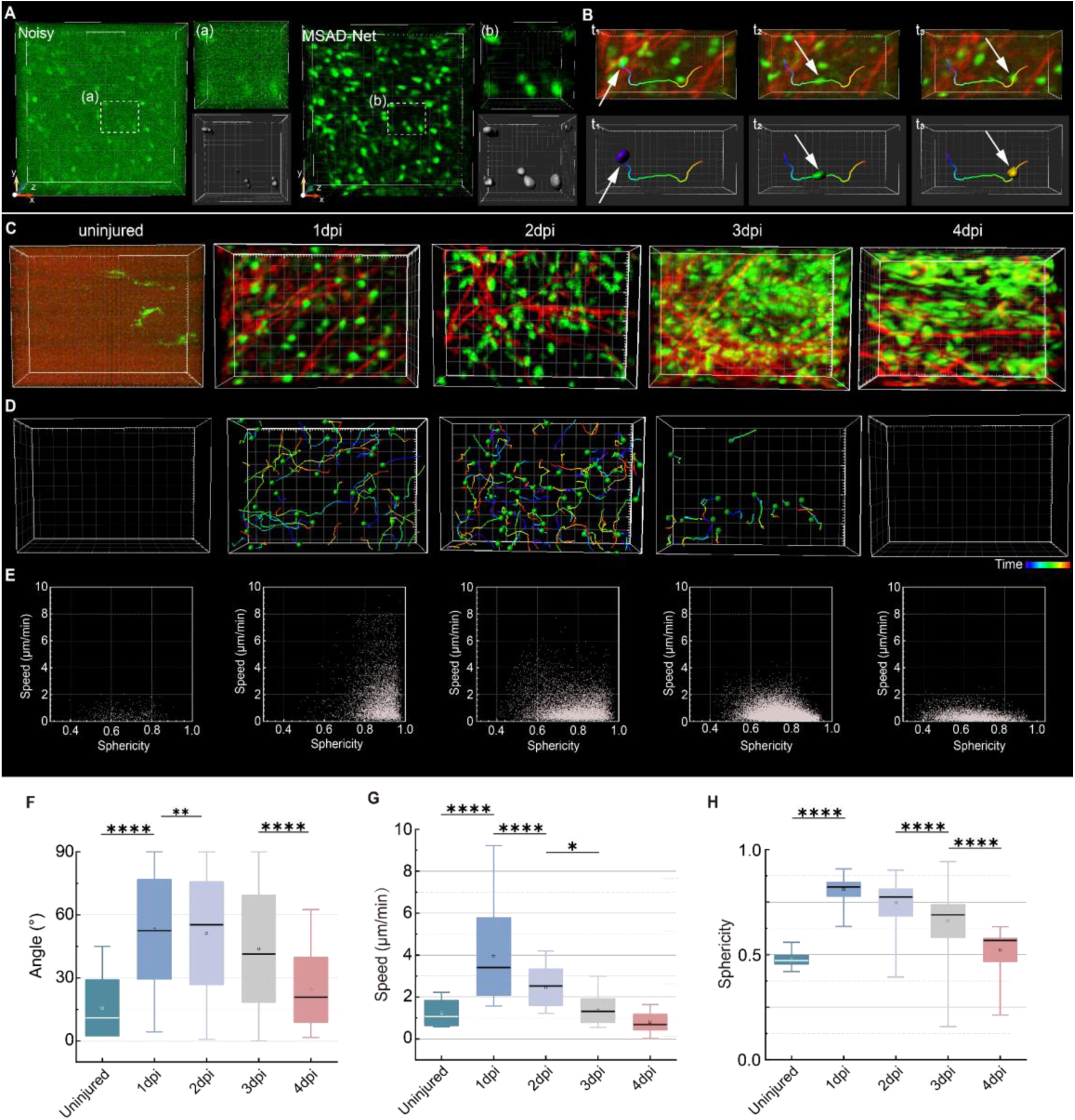
Dynamic tracking of macrophages during muscle regeneration, via 3D three-photon fluorescence and SHG microscopic images denoised by the MSAD-Net. (A) Noisy and denoised 3D three-photon fluorescence microscopic images of macrophages. The upper right panels showed insets (a) and (b) in the noisy and denoised images, respectively. The bottom right panels displayed the corresponding extracted 3D morphology of macrophages. (B) Process of extracting the 3D motion trajectory of macrophages. The upper panels presented the 3D three-photon fluorescence microscopic images of macrophages from t_1_ to t_3_. The bottom panels showed the corresponding extracted 3D morphology of macrophages from t_1_ to t_3_. Arrows indicated the position of macrophages. The colors ranging from purple to red represented the chronological progression. (C) Three-photon fluorescence microscopic images of macrophages (displayed in green) and SHG microscopic images of muscle fibers (displayed in red) at different injury time points of muscle. (D) 3D dynamic trajectory of macrophages at different injury time points based on the trajectory extraction process in (B). (E) Scattering diagram of sphericity and motion velocity of macrophages. The sphericity and motion velocity were calculated according to the morphology and trajectory of macrophages. Details were described in ‘Methods’ part. (F) Variation in the angle between the long axis of macrophages and the SHG signal from muscle fibers with injury time. (G) Variation in motion velocity of macrophages with injury time. (H) Variation in sphericity of macrophages with injury time. Data in (F-H) were analyzed using one-way ANOVA with Tukey’s multiple comparison test. Error bars represent the standard deviation. *P < 0.05; **P < 0.01; ***P < 0.001; ****P < 0.0001.

We performed morphological and trajectory analysis of macrophages at different time points (Fig. 5C-E, Supplementary Video 3-7). Under non-injury conditions, macrophages were sparsely present and mostly stationary, distributed along the muscle fibers. However, at 1 dpi and 2 dpi, macrophages accumulated at the injury site with increased average angle between the macrophages’ orientation and SHG signal from muscle fibers (Fig. 5F). At 3 dpi, the movement trajectories of macrophages decreased, and the majority of cells returned to a nearly stationary state similar to the non-injured muscle. At 4 dpi, the macrophages stopped moving, and the average angle between the macrophages’ orientation and SHG decreased (Fig. 5F, Fig. 5G).

Interestingly, following muscle injury, macrophages primarily exhibited a spherical morphology with faster movement, which is distinct from the non-injured stage (Fig. 5G-H). As regeneration proceeded, the number of round-shaped macrophages gradually decreased, accompanied by a decrease in movement speed (Fig. 5G-H). Macrophages are a heterogeneous cell population, and different subgroups dominate at different stages of regeneration, exerting corresponding functions. Our results propose a new strategy for defining cell subpopulation based on cell morphology and migration. We classified the round-shaped macrophages as “responsive cells”, which primarily respond to injury by proliferating and guiding fiber orientation, characterized by high-speed movement and spherical morphology. The cells with a lower proportion of round-shaped morphology were classified as “resting cells”, which mainly promote myofiber formation around the fibers and exhibit a stationary characteristic.

In summary, our real-time *in vivo* 3D tracking method uncovered two subtypes of macrophages with distinct morphology and motion velocity, which may correspond to the polarization transition of macrophages during the regeneration.

## DISCUSSION

During imaging of high-scattering biological tissues, achieving high axial resolution is crucial for distinguishing and identifying different cell population along the imaging depth, thereby ensuring accurate downstream quantitative analysis. Our results (Fig. 1) demonstrated that 2PM has limited axial resolution in muscle imaging, while 3PM based on a higher nonlinear optical effect offers better imaging depth and axial resolution. However, 3PM demands high-peak-power excitation laser and longer scanning time, which increases the risk of photodamage to biological tissue. Though the enhancement of the axial resolution of 2PM can be achieved through a supervised approach by obtaining corresponding pairs of 2PM and 3PM images, this method requires very expensive hardware, including both 2PM and 3PM systems. In the current work, we have developed MSAD-Net, a deep learning-based approach, which utilizes lower excitation power and shorter scanning time without causing photodamage to biological tissue and compromising image quality. Our proposed deep learning method can be seamlessly implemented in the general 3PM system in the absence of complex data acquisition or hardware configurations, making it particularly appealing for researchers. While our approach shares similarities with recent supervised learning methods for image restoration, we emphasize its significant advantage of high fidelity compared to previous techniques. This guarantees the faithfulness and precision of the denoised images, effectively reducing spurious structures and preserving tissue integrity. It is particularly useful for downstream analysis, such as morphology and dynamics tracking. Additionally, our network enables real-time denoising of images with diverse structures and varying levels of noise, highlighting the robustness and practicality.

By employing MSAD-Net-assisted 3PM, we have shown the dynamic interactions between MuSCs, macrophages and muscle blood vessels during muscle regeneration. Moreover, the application of 3PM in investigating other vulnerable and high scattering organs, such as the kidney and liver, provides enhanced penetration depth and improved axial resolution capabilities. However, addressing potential photodamage caused by thermal effects and nonlinear plasma effects remains challenging. Our proposed method holds the potential for unraveling light-sensitive and complex biological processes, including the dynamics and interactions during acute and chronic injuries in the kidney, liver, lung, and intestinal inflammation.

The future goal of 3PM is to achieve faster, deeper and wider-field imaging. However, the scanning speed in deep 3PM is typically limited to a few microseconds per pixel due to constraints related to the pulsed laser excitation with relatively low repetition rate, which affects imaging speed and field of view. The SBR of images at large depths is often low. Our proposed method has solved the issue of limited scanning speed in 3PM to some extent. The future application of deep learning methods holds the promise to address other challenges. For example, self-supervised deep learning methods can also enhance the SBR of images obtained at large depths by learning from high-SBR images at shallow depths. Moreover, training on high-resolution images acquired under objectives with high numerical aperture (NA) enables large-scale imaging while maintaining high resolution. To summarize, various deep learning methods can be adopted by biological laboratories to improve the performance of 3PM, enabling the investigation of complex in vivo biological processes in high scattering tissues with high-resolution large-depth 3D imaging.

## METHODS

### Materials

The Pax7CreERT2/+ mice were provided by Charles Keller (Oregon Health & Science University, Portland, Oregon). The H11-CAG-tdTomato mice (Strain NO. T006772) were purchased from GemPharmatech (Nanjing, China). The KDR-GFP mice (vascular endothelial cells were labeled by GFP) were provided by Center for Genetic Medicine, The Fourth Affiliated Hospital, Zhejiang University School of Medicine. The mice were of the C57BL/6 background, and their genotype was determined by PCR using tail DNA. They were housed in a specific-pathogen-free animal care facility at Zhejiang University, where they had free access to water and standard rodent chow. CX3CR1-GFP mice (macrophages were labeled by GFP) were provided by the Britton Chance Center for Biomedical Photonics, Wuhan National Laboratory for Optoelectronics - Advanced Biomedical Imaging Facility, Huazhong University of Science and Technology. The mice were housed in cages with two to five animals each, following a standard 12-hour light/dark cycle, in a temperature-controlled environment (22 to 25°C) with 40% to 60% humidity. They were provided ad libitum access to food and water. All animal experiments and housing procedures were conducted in accordance with the guidelines of the Institutional Ethical Committee of Animal Experimentation at Zhejiang University (ZJU20230386).

### Tamoxifen injection and BaCl_2_-induced muscle injury

Tamoxifen (T5648, Sigma-Aldrich) was dissolved in sunflower seed oil and was intraperitoneally injected into 2-month-old mice at 12 mg per 100 g of body weight for 5 consecutive days (0d-5d). Then, at the 6th day (6d), 50 µl of 1.2% BaCl_2_ (342920, Sigma-Aldrich) dissolved in saline was injected into the tibialis anterior muscles of mice after tamoxifen withdrawal to induce muscle injury (0dpi). Imaging was conducted at 0-4 day(s) post injury (0dpi-4dpi).

### Preparation for muscle tissue imaging

The mice were anesthetized with pentobarbital sodium before surgery. The skin covering the TA muscle was resected to expose approximately 0.5 cm^2^ of the muscle. The fascia and epimysium were gently removed to avoid damaging the muscle tissue. Then, a glass slide was delicately placed on the muscle tissue, taking care not to apply pressure. The glass slide was secured onto a stand for imaging. A temperature-controlled heating pad set at 35°C was placed beneath the mouse to maintain its body temperature.

### Nonlinear optical microscopic imaging of mouse muscle

In Fig. 1A-H, three-photon and two-photon fluorescence imaging of vascular endothelial cells and MuSCs was performed using the Ultima *in vivo* scanning microscopic system (Bruker Corporation) with a wavelength-tunable femtosecond (fs) laser (Spectra-Physics, Spirit & NOPA-VISIR) as the excitation source. For imaging vascular endothelial cells, the 850 nm fs laser beam was selected for two-photon fluorescence excitation, while the 1300 nm fs laser beam was used for three-photon fluorescence excitation. To image MuSCs, the 1150 nm fs laser beam was used for two-photon fluorescence excitation, and the 1550 nm fs beam was selected for three-photon fluorescence excitation. The commercial Bruker microscopic system was equipped with an 800 nm long-pass dichroic mirror and a water-immersion objective (XLPLN25XSVMP2-SP1700, 25×, NA=1.05, Olympus). The imaging step size was set to 2 μm, and images were captured with a resolution of 512×512 pixels and an average of 2 frames, using an exposure time of 6 μs/pixel. The laser power was adjusted to 5.2 mW for imaging at the surface of muscle tissue and 60 mW for imaging at the maximum depth (1200 μm for vascular endothelial cells imaging and 350 μm for MuSCs imaging). Imaris 9.0.1 software was used for extracting cell morphologies, and Gaussian fitting was applied to analyze the axial resolution of vascular endothelial cells and MuSCs.

To repeatedly image the vascular endothelial cells (in Fig. 1I), the fs excitation wavelength was set as 1300 nm. Images were taken with 512×512 pixels and 2 frames average, using an exposure time of 6 μs/pixel. The fs laser power was set as 5.2 mW for imaging at surface of muscle tissue, and 60 mW for imaging at the largest depth (400 μm). In Fig. 1L, the fs laser power was set as 5.2 mW, and imaging depth was 60 μm.

In Fig. 3, the injury model of muscle tissues was carried out by administering injection of BaCl_2_. To conduct multi-channel microscopic imaging of muscle tissues, the following steps was carried out. Initially, we selected the 1300 nm fs laser as excitation. The three-photon fluorescence of macrophages was captured using a 505-545 nm filter while the second harmonic generation (SHG) signal of fibers was collected using a 650-700 nm filter. Then, the fs excitation wavelength was changed to 1550 nm. The SHG signal of fibers was collected using a 760-900 nm filter while the three-photon fluorescence of MuSCs was collected using a 605-625 nm filter. Subsequently, we replaced the 605-625 nm filter by a 505-545 nm filter to collected the third harmonic generation (THG) signal of fiber membranes, while SHG channel was maintained to check the field of view. Finally, we replaced the 505-545 nm filter by a 660-700 nm filter, and intravenously injected DCBT NPs into the mouse. The three-photon fluorescence of blood vessel (filled with DCBT NPs) and SHG signal of fibers were collected under the 1550 nm fs excitation. The maximum powers were set as 20 mW for 1300 nm excitation and 15 mW for 1550 nm excitation to avoid photodamage during continuous deep imaging. The signals from all five channels (3 three-photon fluorescence channels, 1 SHG channel and 1 THG channel) were then combined. Imaris software was employed to extract the morphology of the MuSCs and blood vessel using “Surface” functionality, and the location of macrophages using “Spots” functionality. The five-channel microscopic imaging was performed on uninjured mice and mice at 1 to 4 day(s) post-injury.

In Fig. 4, to perform dual-channel three-photon fluorescence microscopic imaging of vascular endothelium cells and blood vessels (blood microcirculation), DCBT NPs were intravenously injected into the KDR-GFP mice to fill the blood vessels. Under the 1300 nm fs excitation, three-photon fluorescence of DCBT NPs was collected in the 550-580 nm range and three-photon fluorescence of vascular endothelium cells was collected in the 490-510 nm range. Then, high-resolution dual-channel imaging was obtained under 4× zoom of a 25× objective view. Subsequently, repeated dual-channel imaging of vascular endothelium cells and blood vessels was conducted. For uninjured mice and mice at 2 to 4 days post injury, the power of 1300 nm fs excitation laser at the surface of muscle was set at 1.3 mW, with the maximum power at 20 mW. The acquired images were then denoised by the MSAD-Net for subsequent analysis. In fig. S17, the velocity of the targeted vessel was measured using the line scanning mode of the multiphoton fluorescence microscope. A repetitive line was scanned along the central axis of the vessel at a rate of at least 1000 Hz. The line-scanning generated a space-time image, where the three-photon fluorescence signals of DCBT NPs appeared as streaks with a slope equal to the inverse of the speed.

In Fig. 5, time-lapse imaging of macrophages was performed on uninjured mice and mice at 1 to 4 day(s) post-injury. The power of 1300 nm fs excitation laser was less than 1.5 mW, and the scanning time for a single frame image was less than 0.53 s to avoid photobleaching of GFP and photodamage to muscle tissue. Serial optical sections were captured at 2 μm step to a total depth of 60 μm every 2-3 minutes. The scanning field of view of the injured site was 340 μm × 340 μm (512 × 512 pixels). Imaris software was applied to measure the morphology and track the motion of macrophages. The “Spots” functionality was used to extract the location of macrophages and track their trajectory. The “Surface” functionality was used to extract the morphology of macrophages.

### Imaging processing and analysis

To obtain the motion trajectories of macrophages, we employed the “Spots” functionality in Imaris to accurately determine their spatial positions and movements. Manual identification was used as a supplementary approach to enhance accuracy. To assess the sphericity of macrophages, we utilized the “Surface” functionality in Imaris to identify their morphology. Manual identification was also employed as a supplementary approach to ensure precise results.

To determine the angle between various types of cells and the SHG signal from fibers, we first utilized the “Surface” functionality in Imaris to extract cell morphology and identify the orientation of cell’s long axis. Manual identification was employed when necessary. Subsequently, by subtracting the angle between the long axis of cells and the direction of the SHG signal, we obtained the angle between the cell and the SHG signal.

Furthermore, we utilized the “Surface” functionality in Imaris to compute the distances between macrophages and MuSCs, as well as between macrophages and blood vessels. For instance, to calculate the distance between cells A and B, we initially determined the position of cell A. Subsequently, we computed the distance from each individual cell A to its nearest neighbor cell B. Manual identification was used as a supplementary approach to enhance accuracy.

The bounding box algorithm is a technique used to determine the optimal enclosing space for a set of discrete points. It involves approximating complex geometric objects with slightly larger and simpler bounding boxes. Commonly employed bounding box algorithms include Axis-Aligned Bounding Box (AABB), bounding sphere, Oriented Bounding Box (OBB), Fixed Directions Hulls (FDH) and k-DOP. In this study, we specifically utilized the “AABB bounding box” functionality in Imaris to define the length of the bounding box.

### Multi-Scale Attention Denoising Network (MSAD-Net)

Convolutional Neural Networks (CNNs) excel at extracting local features but struggle to capture global dependencies. To overcome this limitation, we propose MSAD-Net, a hybrid architecture that combines Convolutional Block Attention Module (CBAM) and Transformer modules within the UNet network. This integration enhances both local feature refinement and global contextual awareness. The following outlines the detailed steps:

#### (1) Data Collection and Preprocessing

We collected a dataset consisting of approximately two thousand pairs of three-photon fluorescence microscopic images, encompassing various sample types and imaging conditions. During the image preprocessing stage, we applied several operations, including grayscale transformation, histogram equalization, Gaussian filtering and normalization. Additionally, we augmented the dataset and improved the network’s robustness by employing random flipping, rotation, scaling and random cropping.

#### (2) Network Architecture Design

The MSAD-Net architecture consists of three primary components: an encoder, Transformer modules, and a decoder. The encoder extracts hierarchical features from the input noisy image, employing multiple convolutional layers. Each layer is followed by a CBAM module and down sample operations. While the convolutional layers capture local features, the CBAM modules refine feature maps by emphasizing relevant features and suppressing irrelevant ones. Down sample operations reduce spatial dimensions, enabling the capture of larger contexts at deeper levels.

To incorporate global contextual information, we introduce Transformer modules at the neck of the UNet structure. These modules process the down sampled features from the encoder. The feature maps are further down sampled to match the input size of the Transformer. Subsequently, a series of Transformer layers (N = 12) model long-range dependencies and global context. The processed features are then up sampled to match the input size of the decoder.

The decoder’s role is to reconstruct the denoised image from the encoded features. Its structure is symmetrical to the encoder and comprises convolutional layers and up sampling operations. Each convolutional layer refines the up sampled features to enhance details and reduce artifacts, while the up sampling gradually restores the feature maps to the original input size. Skip connections, which combine low-level and high-level features, aid in preserving detailed information.

#### (3) Residual Connections

To enhance feature propagation and restoration capabilities, we introduce residual structures to replace the convolutional modules in the UNet. This modification enables the network to better learn details and texture information present in the images. It alleviates the problem of gradient vanishing, increases network depth, and enhances the network’s noise suppression ability, thereby improving its generalization capability. We construct down sampling and up sampling modules based on residual connections.

The original UNet structure used convolutional kernels with a size of 3. However, smaller kernels struggle to capture larger receptive fields and contextual semantic features, while larger kernels lead to an increase in network model parameters. To address this issue, we set the kernel size to 3 in the first and second down sampling stages, where the feature maps represent low-level semantic features. This reduces the number of network parameters. In the third and fourth down sampling stages, we set the kernel sizes to 5 and 7, respectively, as the output feature maps contain high-level semantic features. Increasing the kernel size enlarges the receptive field of each convolution operation, enabling the extraction of more high-level semantic features in a single convolution. For down sampling, we employ max pooling as it can extract the most representative and salient features from the images, preserving stripe details in denoising tasks more effectively than average pooling.

#### (4) Attention Mechanism

In the original UNet, we replaced the skip connections with the CBAM module. The CBAM module consists of spatial attention and channel attention modules. The channel attention module learns correlations between different channels, focusing more on the feature channels that are useful for denoising tasks. The spatial attention module learns correlations between different spatial positions, weighting important regions in the image to enhance the network’s focus on significant structures. This comprehensive approach captures global features in the image, facilitating the transmission of global contextual information between the encoder and decoder.

To enhance the network’s feature extraction capabilities, Transformer modules are introduced at the neck of the UNet structure. These modules, comprising multiple layers (N=12), incorporate multi-head attention mechanisms and feed-forward neural networks. The feature maps from the encoder are down sampled and processed through the Transformer layers, capturing long-range dependencies to enhance global feature representation.

Within each Transformer layer, the attention mechanism follows layer normalization and a multi-layer perceptron (MLP), ensuring stable and effective learning of global features. The enhanced feature maps from the Transformer are then up sampled and passed to the decoder, facilitating the transmission of rich global contextual information.

#### (5) Loss Function and Training

To train the network, we devised a hybrid loss function that combines mean squared error (MSE) and multiscale structural similarity index (MS-SSIM). The MSE loss function quantifies the discrepancy between the network’s output image and the corresponding ground truth image. Additionally, we employed the multiscale structural similarity index loss function, which aids in preserving high-frequency information, including image edges and fine details. By incorporating the multiscale structural similarity index loss function, our network demonstrates improved performance in preserving image stripe details while denoising.

The formula for Mean Squared Error (MSE) loss function is as follows:

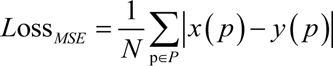

where *N* is the number of samples, *x(p)* represents the true value of the p-th sample, and *y(p)* represents the model’s predicted value for the p-th sample.

The formula for the Multiscale Structural Similarity Index (MS-SSIM) loss function is as follows:

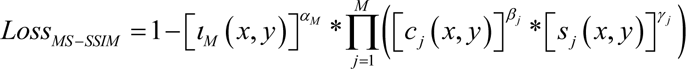

In the equation, *l(x, y)* represents the mean brightness of an image, *c(x, y)* represents the variance of brightness, which is used to estimate contrast, and *s(x, y)* represents the covariance of brightness, which is used to estimate structural similarity. *M* represents the number of scales, which is set to 5. This means that there are a total of 5 scales, including 4 down sampled scales and the original scale. *α_m_* denotes the importance weight for luminance correlation at the last scale. *β_j_*represents the importance weight for contrast correlation at the j-th scale. *γ_j_* represents the importance weight for structure correlation at the j-th scale. In the experiment, these parameters are set as follows:

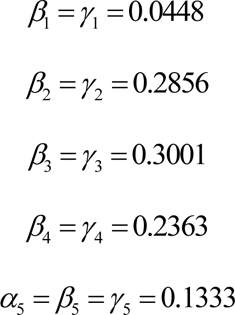

The final loss function is as follows:

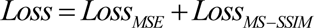

We used the Adam optimizer for training the network, with an initial learning rate of 0.001. The learning rate scheduler was set to Cosine Annealing Warm Restarts.

The network was implemented using PyTorch and executed on a PC equipped with an Intel Core i7-7700HQ CPU @ 2.80GHz, 8 GB RAM, and a Nvidia GeForce GTX 1070 GPU. By employing the above-mentioned methods, we successfully implemented the denoising task for three-photon fluorescence and other nonlinear optical microscopic images using an improved UNet network. The experimental results demonstrate significant improvements in reducing noise levels, preserving image details, and enhancing visual quality. These achievements hold significant application value for image analysis and subsequent processing tasks.

### Comparison Methods

We compared the denoising performance of our optimized architectures with several methods, including Median Filtering, Gaussian Filtering, BM3D, Pix2pix, DNCNN, Masked Denoising, and UNet. Below, we provide implementation details for these methods.

Median filtering and Gaussian filtering were implemented using Fiji. The BM3D method was implemented using MATLAB, and the code is available at https://webpages.tuni.fi/foi/GCF-BM3D/. The noise standard deviation was set to 0.5.

Pix2pix was implemented using the code provided at https://github.com/junyanz/pytorch-CycleGAN-and-pix2pix. The network was trained using the same data as ours, and the default parameters provided in the code were used for training.

DNCNN was implemented using the code provided at https://github.com/SaoYan/DnCNN-PyTorch. The network was trained using the default parameters provided in the code.

Masked Denoising was implemented using the code provided at https://github.com/haoyuc/MaskedDenoising. The network was trained using the default parameters provided in the code.

UNet was implemented using the code provided at https://github.com/milesial/Pytorch-UNet. The network was trained using the same data as ours, and the default parameters provided in the code were used for training.

### Statistical analysis

All imaging data were analyzed using the Image J software (Fiji edition). 3D image reconstruction was performed on the Imaris (Bitplane). Statistical analysis and data visualization were performed using GraphPad Prism 7 and Origin 2021 software. All similar measurements were performed on more than two mice. For comparison between two groups or multiple groups with two samples, unpaired two-tailed t test or one-way analysis of variance (ANOVA) with Tukey’s multiple comparisons test was used.

## Supplementary Materials

This PDF file includes:

Fig. S1 to S17

Table. S1

Legends for movies S1 to S7

References

Other Supplementary Material for this manuscript includes the following:

Movies S1 to S7

## Funding

This work was supported by the National Key R&D Program of China (2022YFB3206000), Dr. Li Dak Sum & Yip Yio Chin Development Fund for Regenerative Medicine, Zheiiang University, National Natural Science Foundation of China (61975172), and Postdoctoral Fellowship Program of CPSF under Grant Number GZB20240646.

The authors acknowledge Shuangshuang Liu from the Core Facilities, Zhejiang University School of Medicine, for her technical support and Prof. Dongyu Li from Huazhong University of Science and Technology, for his constructive suggestion.

## Author contributions

Y.L. was involved in the conceptualization, experimental design, investigation, statistical analysis, writing and editing of the manuscript. K.L. was involved in the experiments, investigation and statistical analysis. M.H. was involved in the experiments, investigation, writing and editing of the manuscript. C.L. and R.Z. were involved in designing deep-learning methods and editing the manuscript. W.X. was involved in investigation and experimental design. S.Q. was involved in providing CX3CR1-GFP mice. Z.W. and M. J were involved in providing KDR-GFP mice. Z.Z. was involved in providing DCBT NPs. X.X. and J.Q. were involved in conceptualization, experimental design and editing of the manuscript.

## Data and materials availability

All data needed to evaluate the conclusions in the paper are present in the paper and/or the Supplementary Material.

