## Supplemental figs for "Ultra-low photodamage three-photon microscopy assisted by neural network for monitoring regenerative myogenesis"

Yifei Li [1], Keying Li [2], Mubin He [1], Chenlin Liang [3], Xin Xie [2], Jun Qian [1]
[1]State Key Laboratory of Extreme Photonics and Instrumentation, Centre for Optical and Electromagnetic Research, College of Optical Science and Engineering, International Research Center for Advanced Photonics, Zhejiang University, Hangzhou 310058, China.
[2]Zhejiang University-University of Edinburgh Institute, Zhejiang University School of Medicine, Zhejiang University, Haining 314400, China.
[3]Tandon School of Engineering, New York University, New York, 11201, the United States.

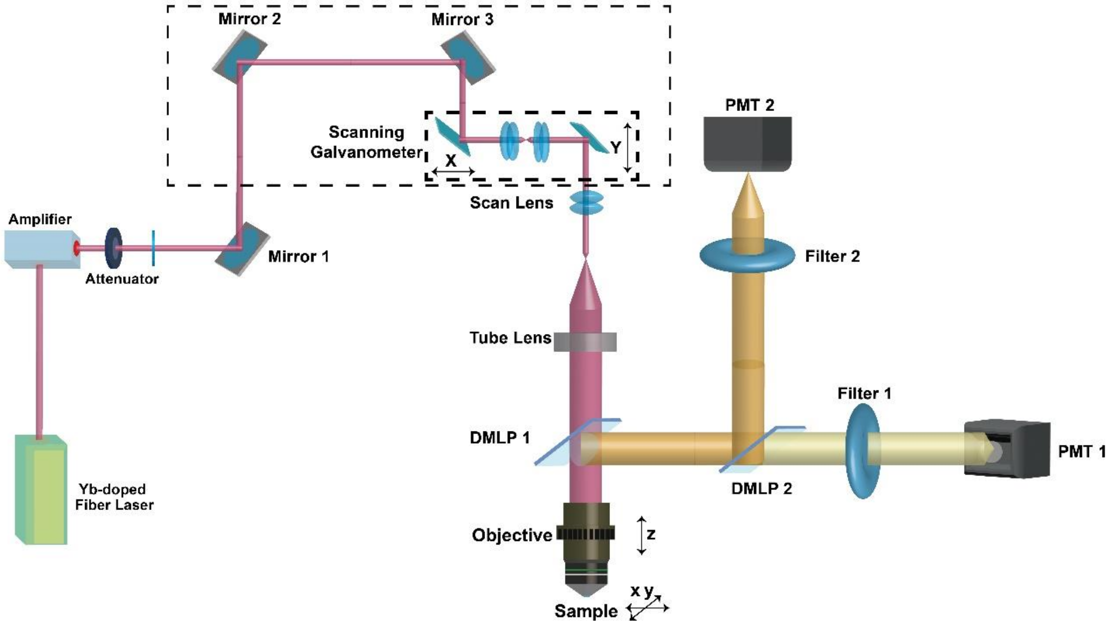


**fig. S1. Schematic illustration of the multiphoton fluorescence microscopic system.**

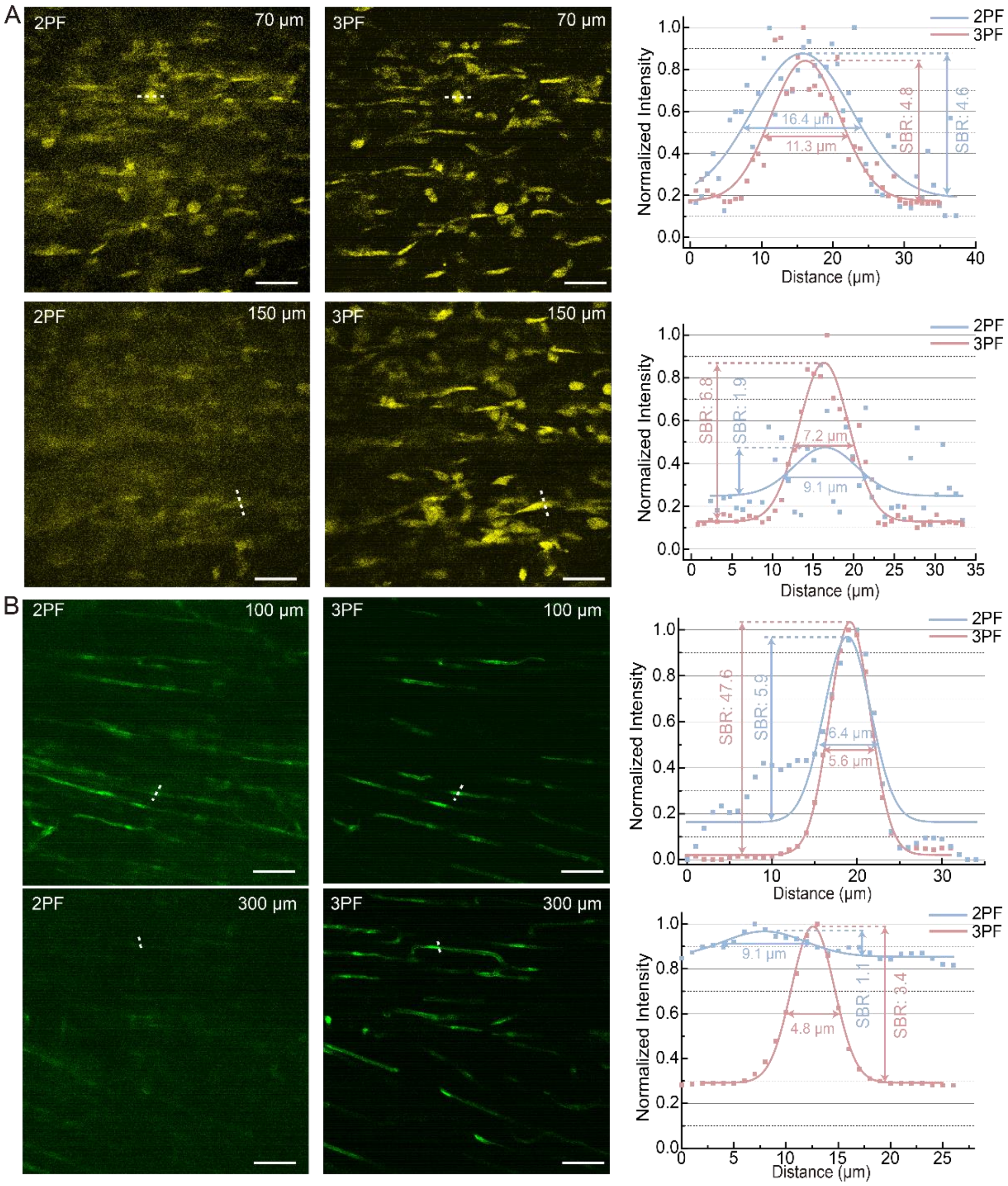


**fig. S2. Three-photon and two-photon fluorescence microscopic images of MuSCs and vascular endothelial cells.** (A) Three-photon and two-photon fluorescence microscopic images of MuSCs at 75 and 150 μm depths. Right panels showed intensity along dotted lines in two-photon and three-photon images in left panels. The values of FWHM (full width at half maximum) and SBR (signal-to-background ratio) indicated in the figures were calculated by gaussian fitting function according to the intensity profiles. Scale bar: 50 μm. (B) Three-photon and two-photon fluorescence microscopic images of vascular endothelial cells at 100 and 300 μm depths. Scale bar: 50 μm. Right panels showed intensity along dotted lines in two-photon and three-photon images in left panels. The values of FWHM and SBR indicated in the figures were calculated by gaussian fitting function according to the intensity profiles. Scale bar: 50 μm.

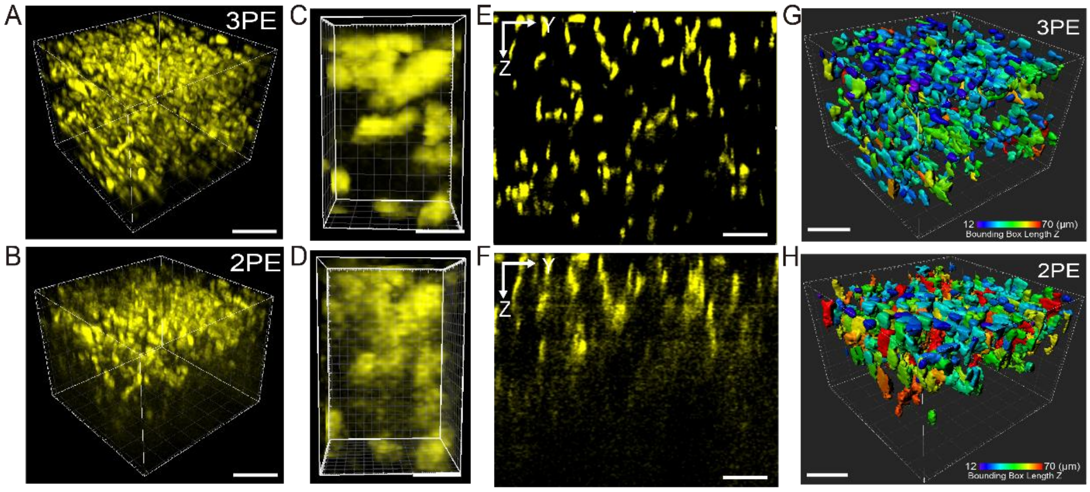


**fig. S3. Three-photon and two-photon fluorescence microscopic images of MuSCs.** (A-B) 3D (A) three-photon and (B) two-photon fluorescence microscopic images of MuSCs. Volume size: 400 μm ×400 μm ×350 μm. Scale bar: 100 μm. (C-D) Enlarged 3D (C) three-photon and (D) two-photon fluorescence microscopic images of MuSCs. Volume size: 30 μm ×30 μm × 50 μm. Scale bar: 10 μm. (E-F) Y-Z plane of (E) three-photon and (F) two-photon fluorescence microscopic images of MuSCs. Scale bar: 50 μm. (G-H) Morphological extraction results of MuSCs from (E) three-photon and (F) two-photon fluorescence microscopic images. The colors ranging from purple to red represented the bounding box lengths in the Z direction.

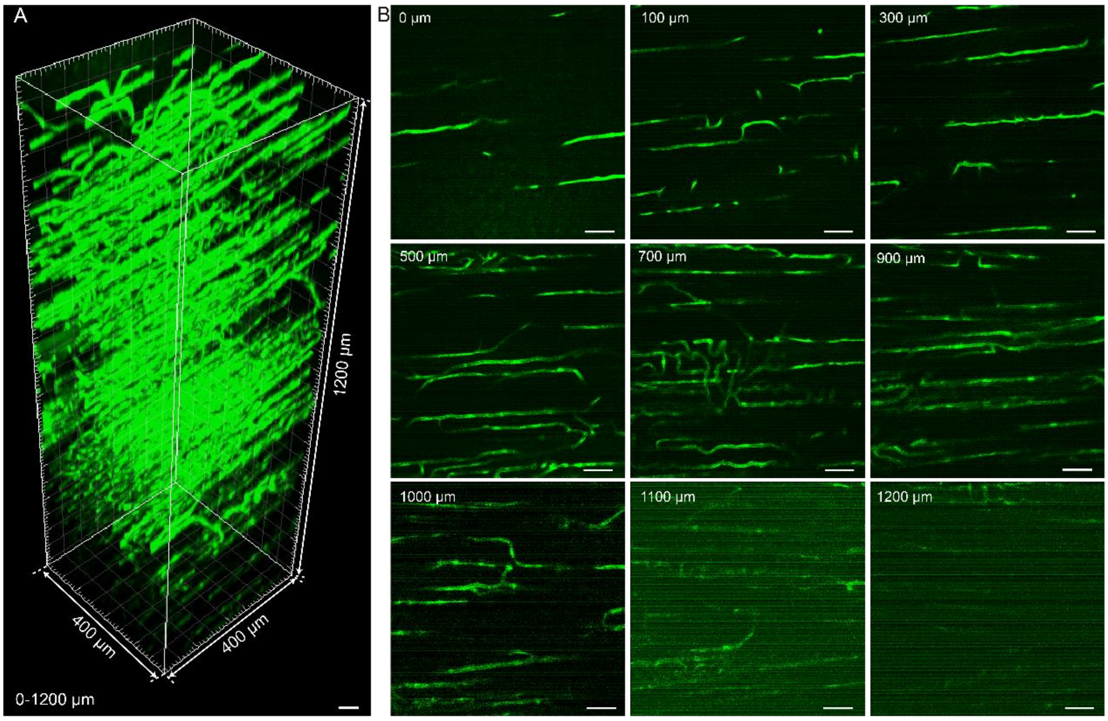


**fig. S4. Three-photon fluorescence microscopic images of vascular endothelial cells.** (A) 3D three-photon fluorescence microscopic imaging of vascular endothelial cells. Volume size: 400 μm ×400 μm ×1200 μm. Scale bar: 50 μm. (B) three-photon fluorescence microscopic imaging of vascular endothelial cells at various depths. Scale bar: 50 μm.

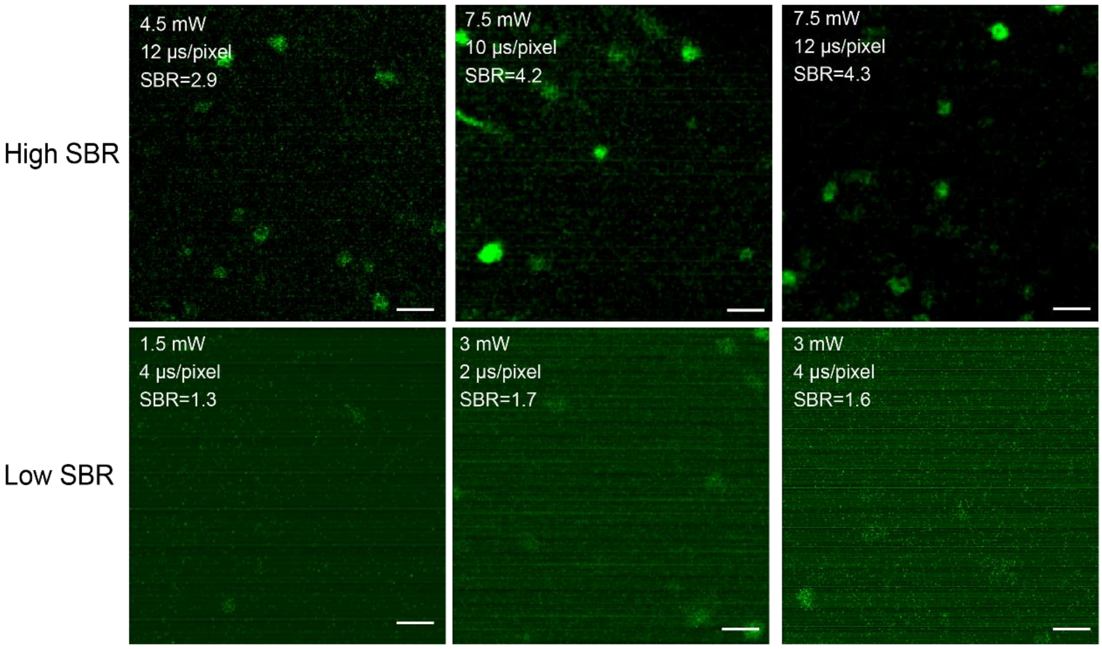


**fig. S5. Three-photon fluorescence microscopic images of macrophage cells with high and low SBRs.** The fs excitation wavelength was set as 1300 nm (1 MHz, 115 fs). The values of SBR indicated in the images were calculated as maximum signal intensity divided by the background intensity. Scale bar: 50 μm.

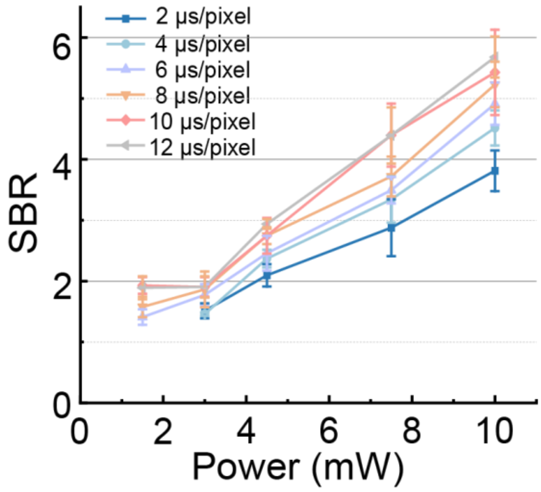


**fig. S6. SBRs of three-photon fluorescence microscopic images of macrophage cells as functions of scanning time and average excitation power at the imaging depth of 50 μm.** The fs excitation wavelength was set as 1300 nm (1 MHz, 115 fs).

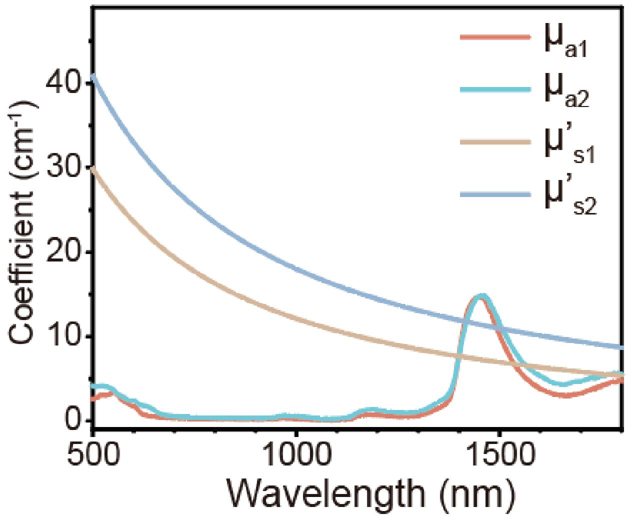


**fig. S7. Absorption and reduced scattering coefficients of muscle tissue.** $\mu_{a1}$ and $\mu_{s1}$ were the absorption and reduced scattering coefficients of muscle tissue before injury, $\mu_{a2}$ and $\mu_{s2}$ were the absorption and reduced scattering coefficients of muscle tissue after injury. Hemispheric reflectance and hemispheric transmittance spectra were measured by an integrating sphere device. Then, absorption and reduced scattering coefficients were calculated from spectra above according to the one-dimensional two-flux Kubelka-Munk (KM) light transport theory.

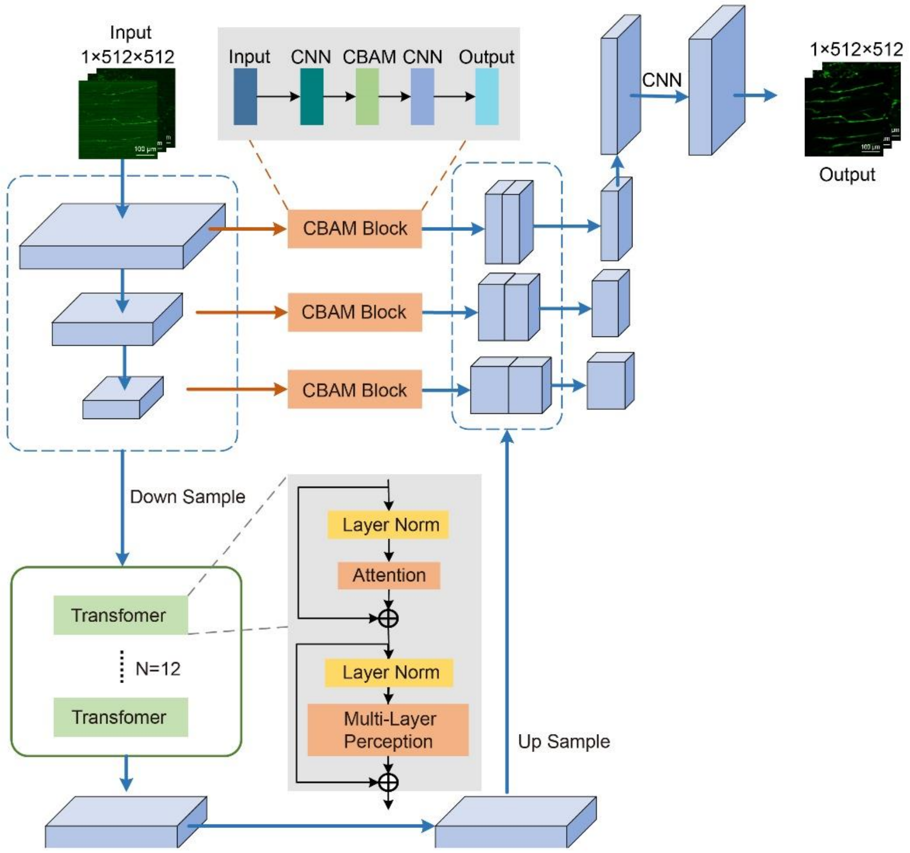


**fig. S8. Schematic of the Multi-Scale Attention Denoising Network (MSAD-Net).**

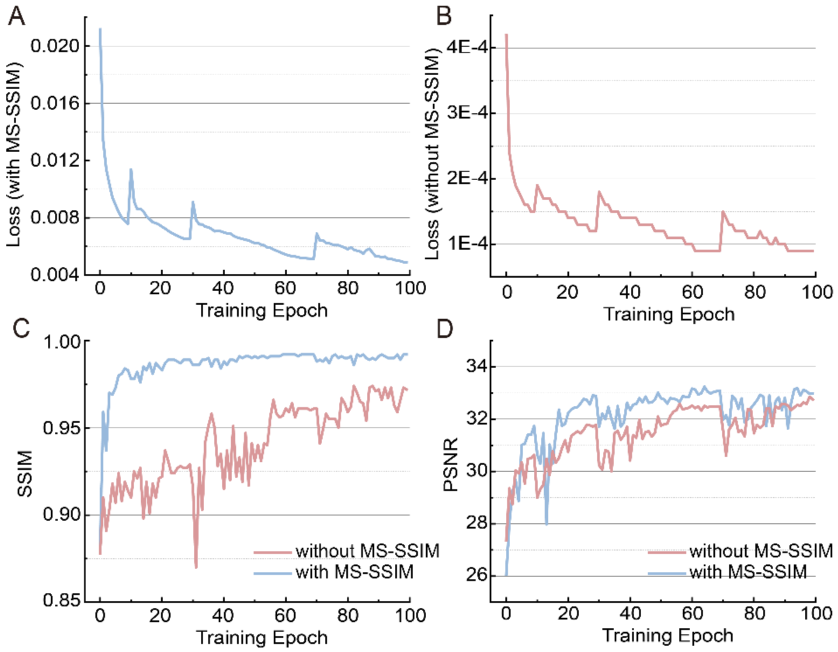


**fig. S9. Several evaluation parameters varied with the number of training epoch.** (a-b) The Loss value using (A) the compound loss function and (B) the single loss function varied with the number of training epoch. (C-D) Variation of (C) SSIM and (D) PSNR with the number of training epoch (using both the single loss function and the compound loss function).

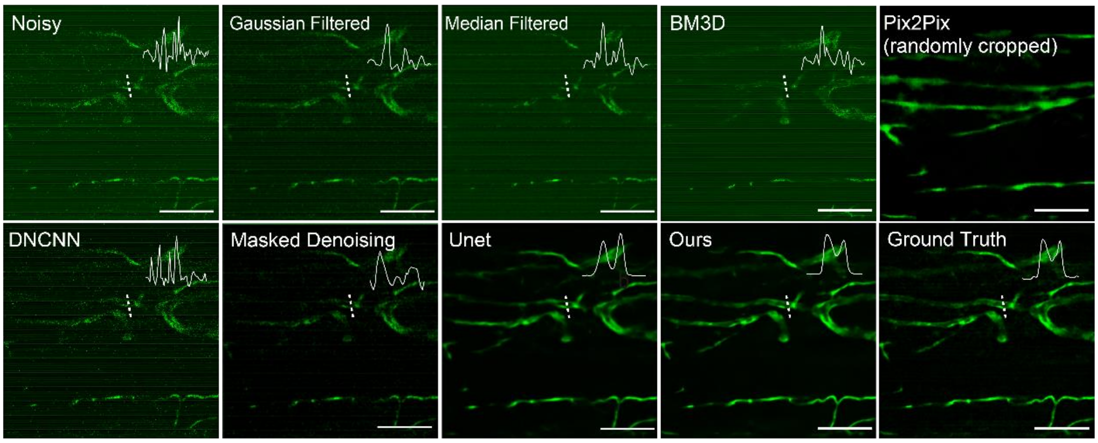


**fig. S10. The original noisy three-photon fluorescence image and corresponding denoised output images of vascular endothelial cells by various methods, including Gaussian filter, Median filter, BM3D, Pix2Pix, DNCNN, Masked Denoising, UNet and MSAD-Net. The three-photon fluorescence image with high SBR was set as Ground Truth for comparison.** Imaging depth was 150 µm. Imaging condition: 6 mW excitation power and 3 µs/pixel scanning time for noisy imaging while 15 mW excitation power and 12 µs/pixel scanning time for Ground Truth imaging. Scale bar: 100 µm.

| Methods | SSIM | PSNR | MSE |
|---|---|---|---|
| Raw images | 0.3040 | 14.4265 | 0.0429 |
| Gaussian filter | 0.2759 | 14.2639 | 0.0318 |
| Median filter | 0.2711 | 14.4314 | 0.0319 |
| BM3D | 0.2820 | 15.6132 | 0.0332 |
| DNCNN | 0.5935 | 22.2557 | 0.0305 |
| Masked Denoising | 0.8313 | 27.2132 | 0.0068 |
| Unet | 0.9684 | 32.3487 | 0.0027 |
| Ours | 0.9932 | 34.6625 | 0.0028 |

**Table. S1. Average values of SSIM, PSNR and MSE comparing the noisy images, and denoised output images by various methods (including Gaussian filter, Median filter, BM3D, DNCNN, Masked Denoising, UNet, and MSAD-Net) with the ground-truth images with high SBRs.**

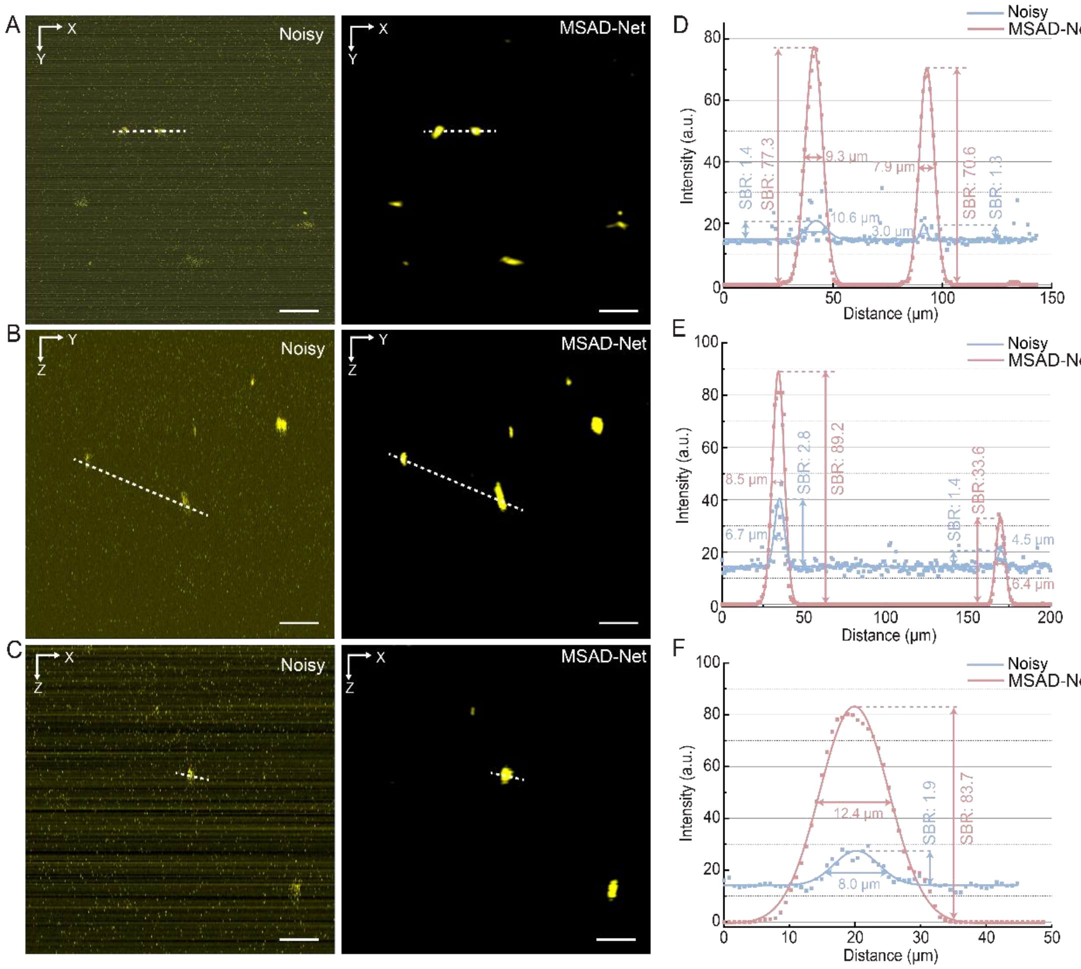


**fig. S11. Three-photon fluorescence microscopic images of MuSCs in uninjured mice, before and after denoised by the MSAD-Net.** (A-C) (A) X-Y plane, (B) Y-Z plane, and (C) X-Z plane images of MuSCs in uninjured mice, before and after denoising by the MSAD-Net. Scale bar: 50 μm. (D-F) Intensity profiles along the white dashed lines across MuSCs in A-C). The values of FWHM and SBR indicated in the figures were calculated using a Gaussian fitting function based on the intensity profiles.

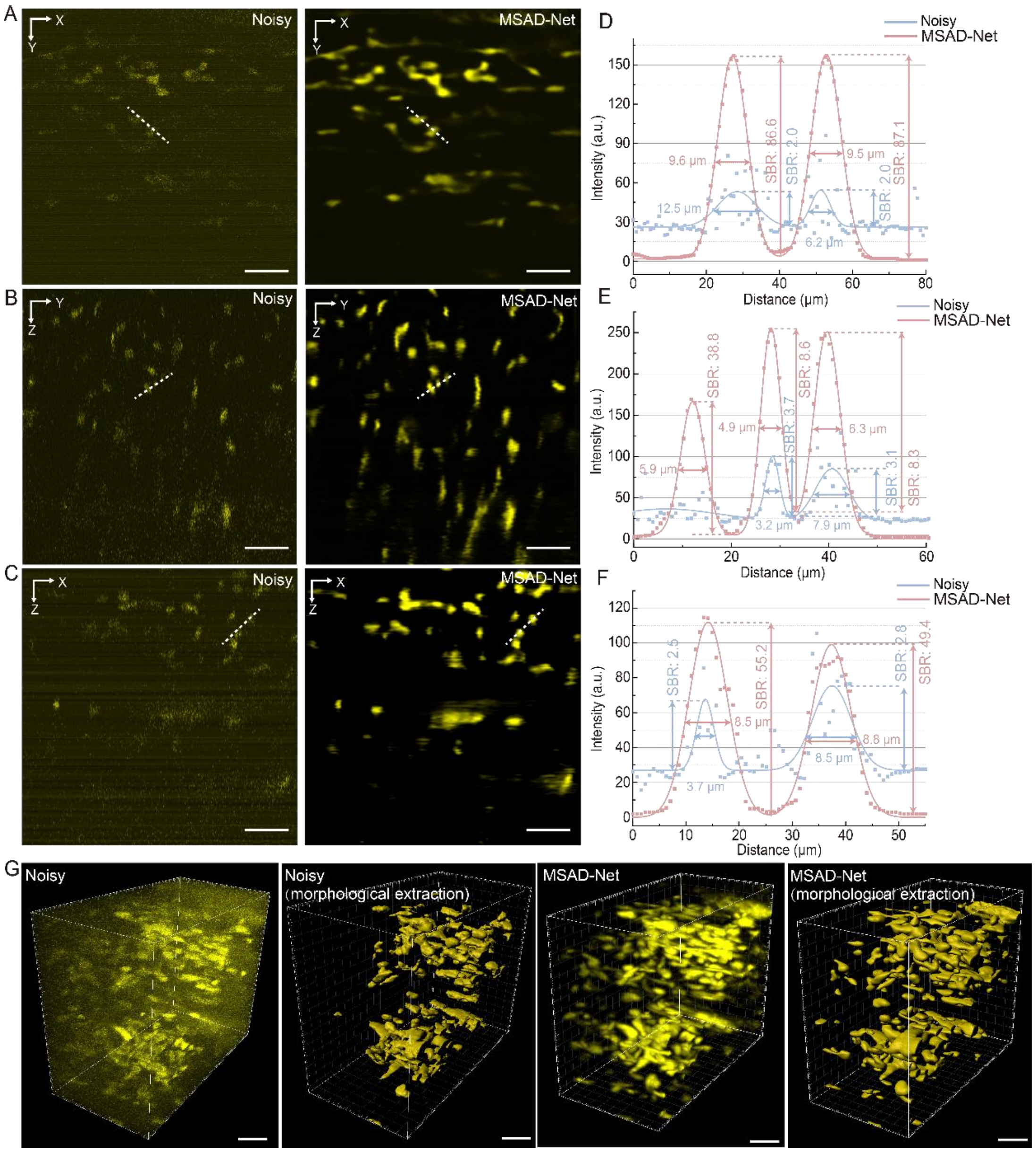


**fig. S12. Three-photon fluorescence microscopic images of MuSCs in injured mice (2dpi), before and after denoised by the MSAD-Net.** (A-C) (A) X-Y plane, (B) Y-Z plane and (C) X-Z plane images of MuSCs in injured mice (2dpi), before and after denoised by the MSAD-Net. Scale bar: 50 μm. (D-F) Intensity profiles along the white dashed lines across MuSCs in (A-C). The values of FWHM and SBR indicated in the figures were calculated using a Gaussian fitting function based on the intensity profiles. (G) 3D reconstruction and morphological extraction results of MuSCs before and after denoised by the MSAD-Net. Volume size: 190 μm ×190 μm ×400 μm. Scale bar: 50 μm.

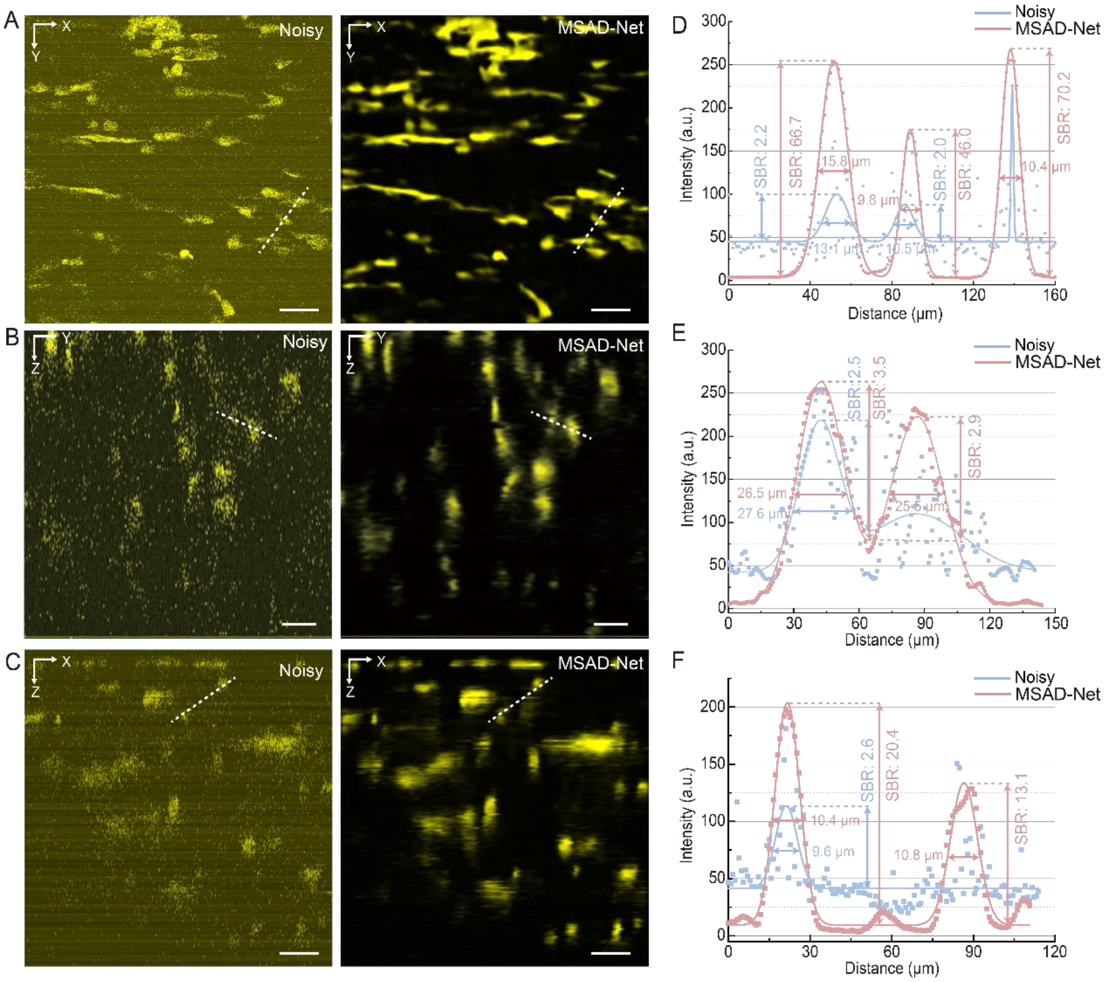


**fig. S13. Three-photon fluorescence microscopic images of MuSCs in injured mice (3dpi), before and after denoised by the MSAD-Net.** (A-C) (A) X-Y plane, (B) Y-Z plane and (C) X-Z plane images of MuSCs in injured mice (3dpi), before and after denoised by the MSAD-Net. Scale bar: 50 μm. (D-F) Intensity profiles along the white dashed lines across MuSCs in (A-C). The values of FWHM and SBR indicated in the figures were calculated using a Gaussian fitting function based on the intensity profiles.

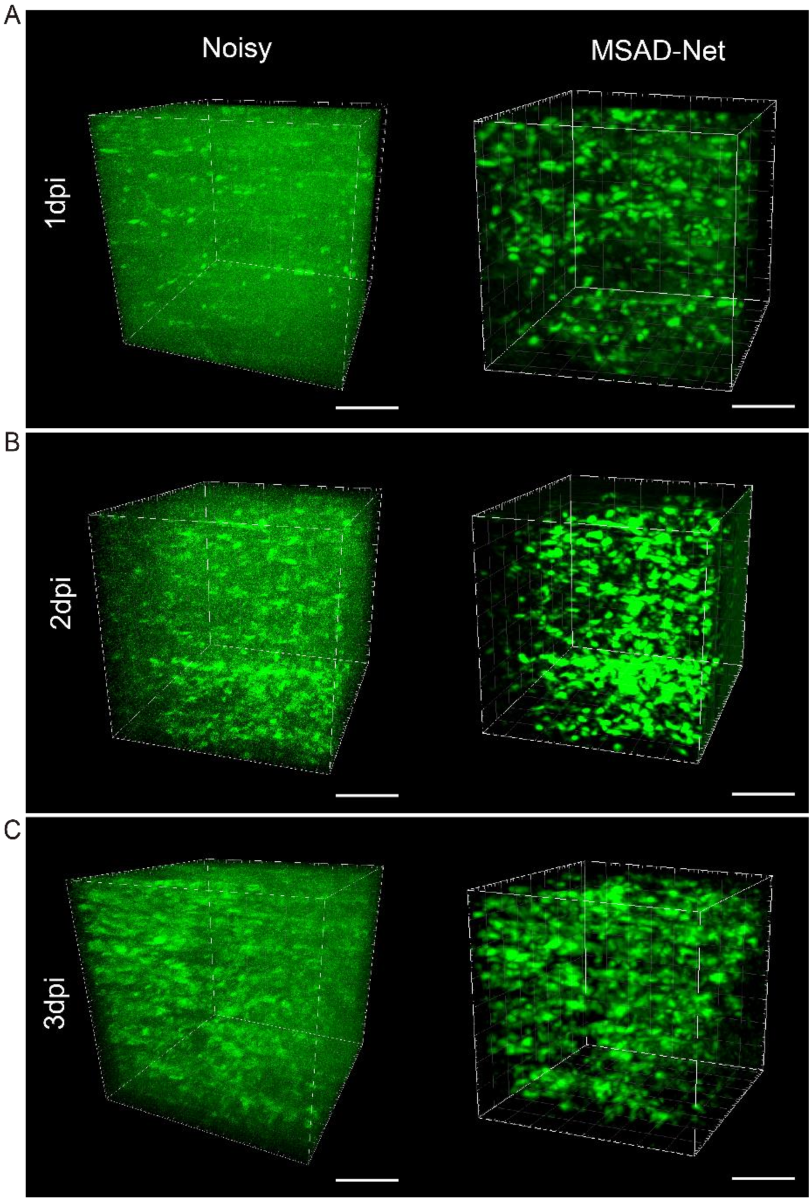


**fig. S14. 3D reconstruction of three-photon fluorescence microscopic images of macrophage cells before and after denoised by the MSAD-Net, in mice at (A) 1dpi, (B) 2dpi mice, and (C) 3dpi.** Volume size: 340 μm ×340 μm ×400 μm. Scale bar: 50 μm.

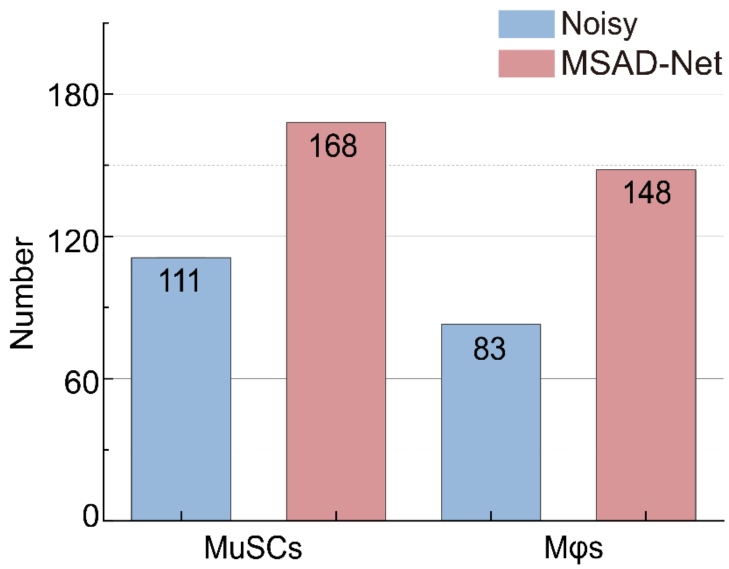


**fig. S15. Extracted number of macrophage cells and MuSCs based on noisy images and denoised images (by the MSAD-Net).**

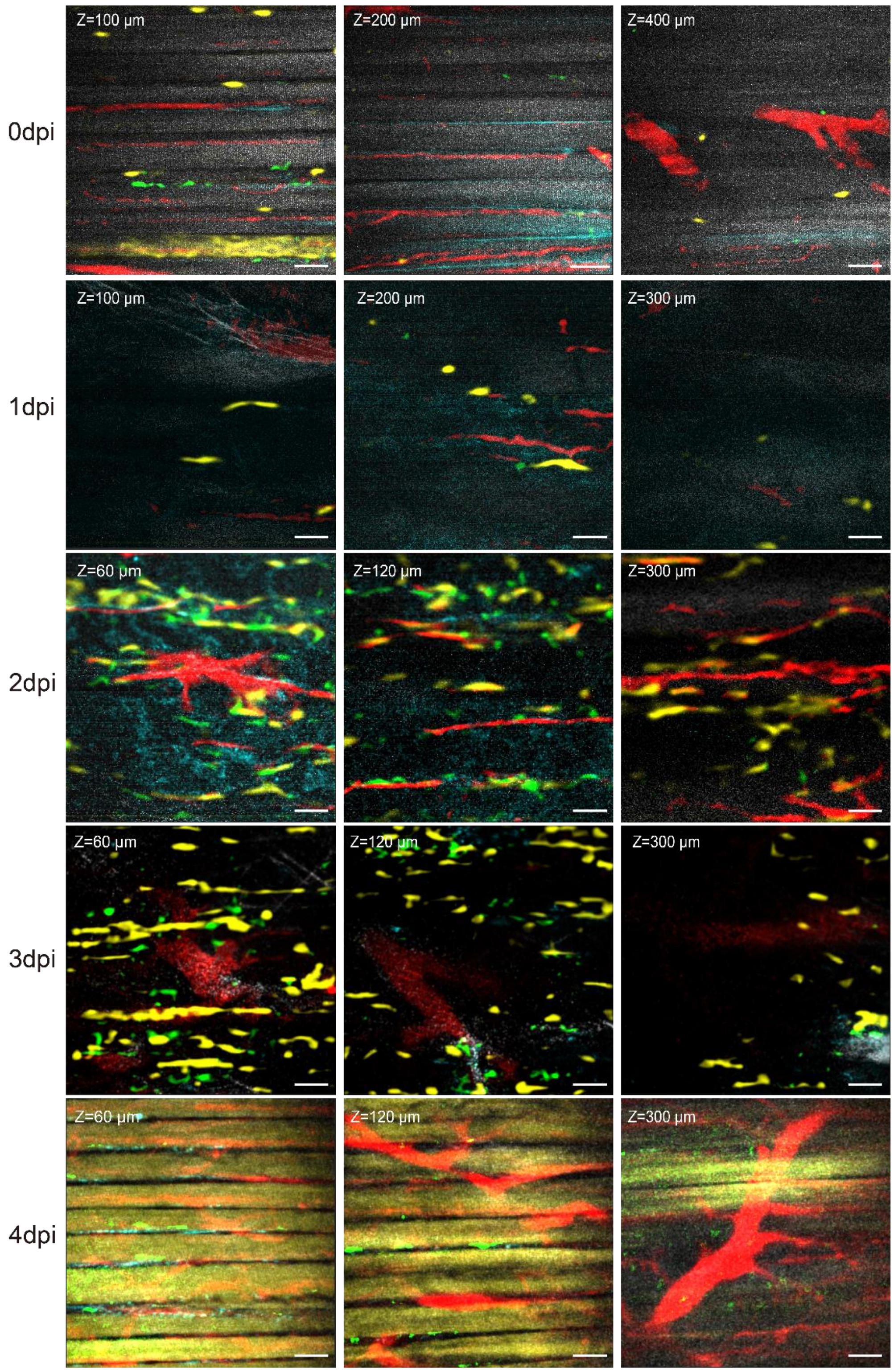


**fig. S16. Five-channel microscopic muscle images, including three-photon fluorescence images of macrophage cells, MuSCs and muscular blood vessels, SHG images of fibers, and THG images of fiber membranes, at various depths in uninjured mice and mice at 1-4 day(s) post-injury (1dpi-4dpi), after denoised by the MSAD-Net.** Scale bar: 50 µm.

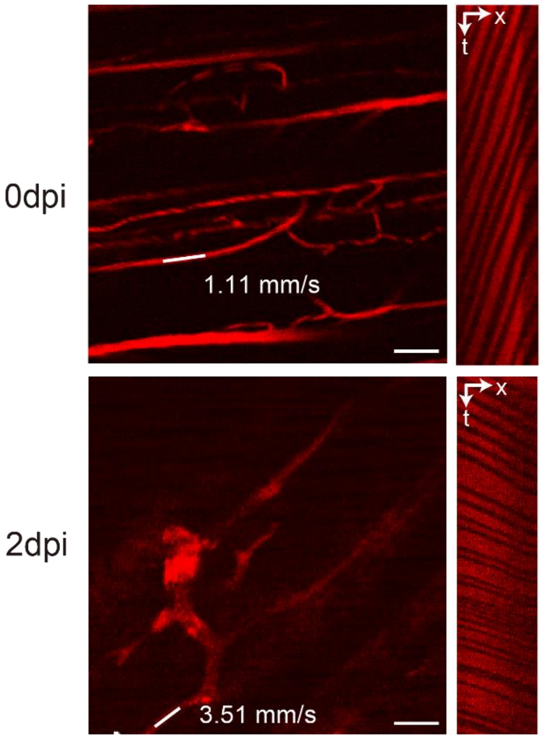


**fig. S17. Measurement of blood flow velocity of uninjured mice and mice at 2 days post-injury (2dpi) base on the three-photon fluorescence microscopy.** Details were described in "Methods" part. Scale bar: 50 μm.

**Supplementary Movies:**

**Movie. S1. Three-photon fluorescence microscopic images of vascular endothelial cells.** Volume size: 400 μm×400 μm×1200 μm. Scale bar: 50 μm.

**Movie. S2. The time-lapse three-photon fluorescence imaging of macrophage cells (displayed in green) and SHG microscopic imaging of muscle fibers (displayed in red).** Movie frames are 3D images of 33 slices spaced 2 μm and captured for 11 times under normal excitation power (6 mW) and prolonged scanning time (6 μs/pixel, average of 2 frames). Volume size: 340 μm×340 μm×64 μm Volume: Scale bar: 50 μm.

**Movie. S3-7. The time-lapse three-photon fluorescence images of macrophage cells (displayed in green) and SHG microscopic images of muscle fibers (displayed in red) at 0dpi-4dpi, denoised by the MSAD-Net.** Movie frames are 3D images of 33 slices spaced 2 μm and captured at 3 minutes interval for 180 minutes. Scale bar: 50 μm.
